# RNA-aware tissue preservation workflows for high-quality spatial transcriptomics

**DOI:** 10.64898/2026.09.07.749899

**Authors:** Paolo Cadinu, Christian A. Reardon-Lochbaum, Hao Zhang, Katerina Kalemaki, Brianna R. Watson, Stelios Smirnakis, Scott B. Snapper, Alan B. Cantor, Jeffrey R. Moffitt

## Abstract

Image-based transcriptomic approaches can define, discover, and chart cell types and states within an array of tissues. However, measurement quality depends on RNA integrity, and the modern tissue preservation toolbox was not designed to protect this highly labile molecule. Here we leverage MERFISH to show that tissue-dependent differences in endogenous RNase activity can shape spatial transcriptomics data quality for different preservation methods and that RNase-activity-guided protocol optimization can improve data quality. In parallel, we introduce an RNA-aware pan-tissue preservation approach, Rapid Inhibition and Permanent Inactivation of Nucleases (RIPIN), that rapidly stabilizes samples with a broad-spectrum RNase inhibitor while permitting slow, chemical inactivation. RIPIN produces high-quality MERFISH measurements in all profiled human and mouse tissues, is compatible with clinical workflows, and is easily integrated with frozen or paraffin sectioning. By highlighting how RNA integrity can be lost during tissue processing, our work may inspire the next generation of RNA-aware histology methods.

## INTRODUCTION

Spatial transcriptomic methods have proven to be transformative tools for the study of a wide variety of tissue and cell biology questions through their ability to quantitatively chart gene expression within intact biological samples^1–8^. RNAs can be mapped in space either through spatial capture- or image-based methods, and while these methods have complementary strengths and weaknesses, image-based approaches, with their single-cell resolution and potential for high sensitivity measurements, offer key advantages for specific biological questions^1–8^. Notable image-based methods include those that generate the fluorescent optical barcodes that identify RNAs using padlock probes^9^, e.g., in situ sequencing^10^, STARmap^11,12^, and Xenium^13,14^, and those based upon single-molecule fluorescence in situ hybridization (smFISH)^15,16^, e.g., MERFISH^17–21^, seqFISH^22,23^, and cosMx^24^. In general, the smFISH-based methods that leverage many tens of FISH probes per target RNA, such as MERFISH or seqFISH, have been shown to be capable of exceedingly high RNA detection efficiencies, in some cases several fold greater than that of single cell RNA sequencing^17–23^. These capabilities provide a degree of data quality that has, in turn, empowered a wide variety of biological discoveries, including cell types and states not detected with other methods^20,25,26^. However, as image-based transcriptomic methods have become more widely adopted and have been applied to a much wider diversity of tissue and sample types, data quality has varied between studies^27–31^ and an understanding of how to consistently leverage these tools to provide data of sufficient quality to define cells with a resolution that matches or exceeds other techniques has not emerged.

The unique challenges associated with spatial transcriptomic sample preparation may be one source of inconsistent data quality. The discovery potential of spatial transcriptomic methods arises, in part, because these methods merge two historically distinct types of information: genome-scale transcript abundance—typically associated with genomic methods—and cell-to-tissue-scale morphology—typically associated with histological methods. However, in merging the strengths of these techniques, spatial transcriptomic methods also combine their challenges. RNA is a naturally labile molecule, degraded both chemically by high temperatures or pH and enzymatically by a vast diversity of RNases that are found both within the sample itself or introduced by the researchers while handling the samples^32–34^. Unsurprisingly, the quality of transcriptomic measurements depends on the preservation of RNA integrity; thus, there has been an effort to develop approaches that stabilize RNAs for both bulk RNA extraction^32,34^ and single-cell characterization^35–39^. These approaches largely focus on lowering the activity of endogenous RNases through lower temperatures, chemical inactivation, or chemical inhibition. For example, arguably the most widely used approach—at least for bulk RNA characterization—is a concentrated solution of ammonium sulfate, which is a potent, broad-spectrum RNase inhibitor^34,40^ that is marketed commercially under the name RNAlater^34^. Importantly, the development of methods to stabilize RNAs through the inhibition and inactivation of endogenous RNases in a wide variety of tissues has been a critical aspect of the success of transcriptomic measurements.

In parallel, tissue histology has its own unique challenges. Tissues are typically imaged as thin slices adhered to glass slides or coverslips, and this process requires tissue preparation methods that can largely preserve the structure of the sample, inactivate it metabolically, and ensure strong adherence to the glass support^41,42^. There is a remarkable degree of diversity in the structural and chemical makeup of different tissues and, accordingly, a wide variety of, often tissue-specific, preservation methods have been developed^43^. For example, chemical fixatives, such as paraformaldehyde (PFA) or its alcohol-stabilized form (formalin), or precipitative fixatives, such as alcohols or heavy ions, are commonly used to preserve cellular structures. However, these fixatives can interfere with downstream molecular profiling or distort cellular structure or tissue morphology. Flash freezing of tissues, by contrast, may better preserve molecular features of the tissue but may leave residual metabolic activity within the tissue^44^. In parallel, tissues are often embedded in a support that facilitates sectioning: common examples include optical cutting temperature (OCT) media for samples that will be sectioned cold or paraffin wax for samples cut at room temperature. Similarly to fixation, embedding methods are also often optimized or tailored to both the tissue and the molecular features of interest.

Importantly, the vast majority of histological approaches were developed prior to our modern genomic era^41,42^. As such, there is a limited understanding of how choices made in the histological preparation of different tissues could affect the integrity of RNA and, thus, spatial transcriptomics data quality. This uncertainty is reflected in the substantial variability in fixation conditions—particularly time and temperature— recommended by major spatial transcriptomic platforms, including Xenium (10X Genomics), CosMx (Bruker), and MERSCOPE (Vizgen). For example, in fresh-frozen tissue preparations, Vizgen recommends post-slicing fixation with PFA for 15 minutes at room temperature or 30 minutes at 47 °C, whereas Bruker suggests 2 hours at 4 °C and 10X Genomics recommends 30 minutes at room temperature^45–47^. Similarly, for fixed tissues embedded in OCT or paraffin, Bruker advises fixation for 18–24 hours at room temperature, while 10X Genomics and Vizgen recommend 16–24 hours at 4 °C^46–48^. Moreover, while each of these commercial vendors notes the possibility that preparation may need to be optimized for individual tissues, only limited guidance on how to do so is provided. As such, it remains unclear how best to prepare tissues for image-based spatial transcriptomic measurements, what mechanisms might shape potential tissue-dependent differences, and what guidelines should be used to efficiently optimize preparation methods for different tissues.

Here we sought to address these questions by using MERFISH to explore how image-based spatial transcriptomics data quality varies with different histological preparation methods in a tissue-dependent fashion. Importantly, we find that MERFISH data quality can vary dramatically between different tissues, depending on the preparation method, and that tissue-dependent endogenous RNase activity is a key determinant of this tissue-dependent data quality. We introduce simple assays to measure RNase activity within histological slices and demonstrate that key steps in multiple common histological preservation protocols can be optimized to improve data quality when guided by these RNase-activity measures. Remarkably, we find that RNase activity is resilient to chemical fixation and long, room-temperature PFA fixations are required to substantially reduce activity in many tissues. Motivated by this observation, we introduce an RNA-aware histological framework. In this framework, sample RNAs are rapidly stabilized with a broad-spectrum nuclease inhibitor to then allow the slower but permanent chemical inactivation of RNases. We term this framework Rapid Inhibition and Permanent Inactivation of Nucleases (RIPIN) and introduce a PFA-compatible RNAlater analog that leverages cesium sulfate to rapidly stabilize RNAs and implement this framework. We show that our cesium sulfate preservation buffer preserves samples as well as RNAlater, is compatible with PFA-fixation, integrates into both cryo- or paraffin-embedding workflows, and produces high-quality MERFISH measurements in a range of mouse and human tissues. Moreover, by separating sample stabilization from chemical inactivation steps that can interfere with other genomic methods for characterizing RNA, RIPIN provides a convenient avenue for prescreening RNA quality or collecting samples for the joint bulk and spatial characterization of samples. More broadly, by illustrating that endogenous RNase activity is a major determinant of image-based transcriptomics data quality, we envision that our work may inspire a new generation of RNA-aware histology methods.

## RESULTS

### Tissue-dependent differences in endogenous RNase activity shapes MERFISH performance in fresh frozen samples

To explore how different approaches to tissue preservation may modulate spatial transcriptomics data quality, we screened the performance of a common image-based spatial transcriptomics technique—multiplexed error robust fluorescence in situ hybridization (MERFISH)^17–20^—across a range of preparation methods. To provide a reproducible set of tissues for this characterization, we selected a range of anatomically and functionally distinct mouse tissues—brain, lung, duodenum, and abdominal skin, and we designed a series of MERFISH libraries targeting 222, 449, and 940 genes, with each covering distinct tissue-specific cell-type markers, complemented with genes within several biological pathways (Methods). We designed these libraries with specific tissue sets in mind and confirmed that the selected gene sets could resolve the expected cellular diversity of each tissue in published single-cell RNA sequencing data^49–55^ (Figure S1).

We started with one of the most common tissue preservation methods used in spatial transcriptomics: the rapid freezing of freshly harvested tissue. In this ‘fresh frozen’ protocol, tissue is rapidly harvested from the organ of interest, embedded within a cryoprotectant—OCT—and then frozen. Thin (∼10 micron) slices are then cryosectioned from the frozen block, rapidly melted onto coverslips, and briefly fixed with PFA to metabolically inactivate the tissue and increase its adherence to the coverslip or slide (Figure 1A). Notably, in some cases, this post-fixation step has been performed at lower temperatures to inhibit enzymatic activity within the slice during this process^47,56^.

**Figure 1.**
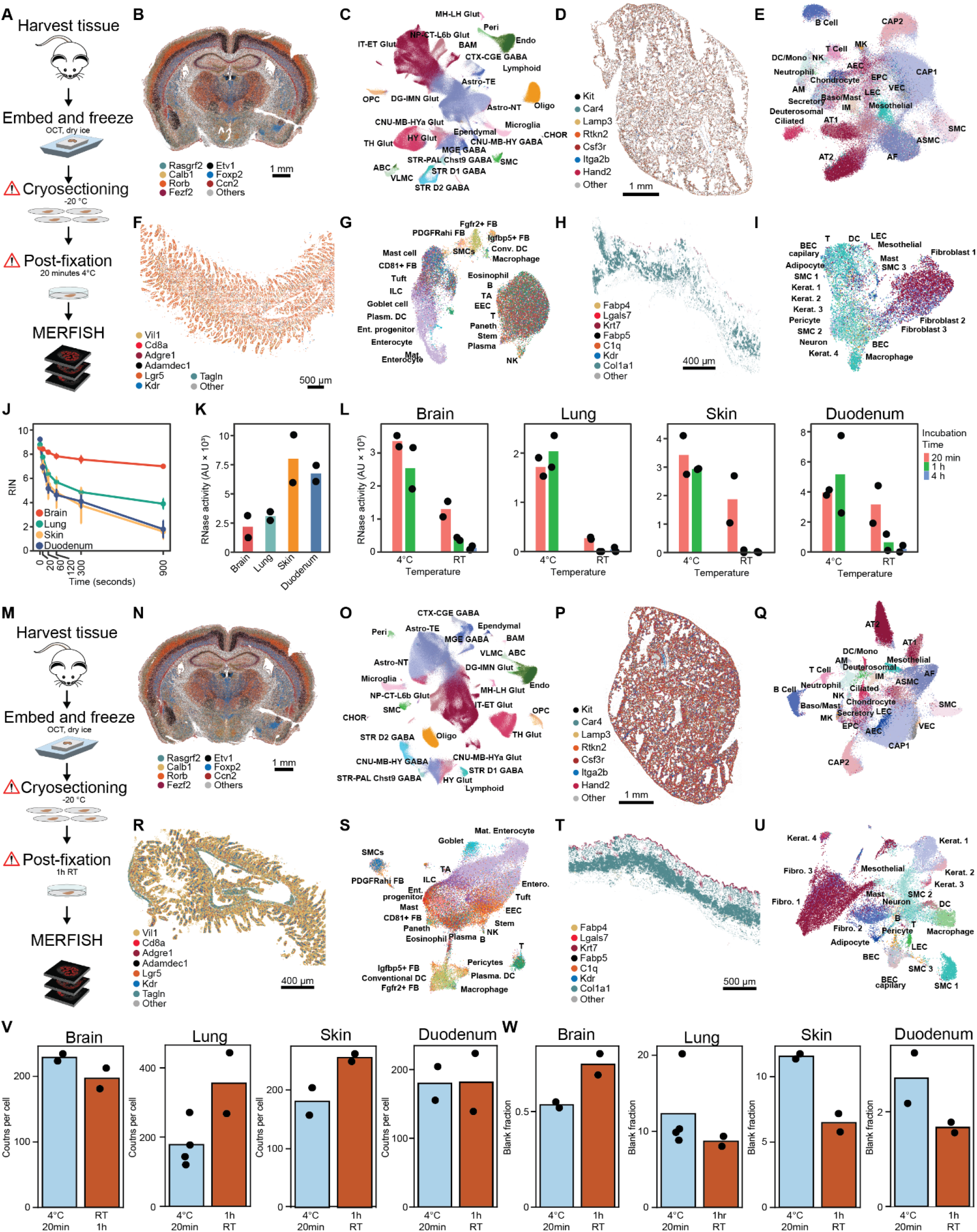
Tissue-dependent differences in endogenous RNase activity shapes MERFISH performance in fresh frozen samples. (A) Schematic workflow for a fresh frozen tissue preparation for MERFISH. (B) Spatial distribution of 7 of 449 mRNAs measured with MERFISH in a coronal section of the mouse brain prepared as in (A). (C) UMAP representation of the cellular diversity seen in the mouse brain with the protocol in (A), colored by major cell type labels transferred to the data from a reference dataset (Methods). (D-I) As in (B,C) but for the mouse lung (D,E), duodenum (F,G), and abdominal skin (H,I). The lung and skin were measured with a 222-gene library while the duodenum was measured with a 940-gene library. (J) RNA integrity number (RIN) for individual tissue slices prepared from fresh frozen tissue blocks as a function of time at room temperature post slicing. Error bars represent standard error of the means across three biological replicates. (K) RNase activity measured by the generation of fluorescence from a test RNA in arbitrary units (Methods) for slices of fresh frozen tissue blocks for the listed tissues. Markers represent replicates while bars represent averages. (L) Per-slice RNase activity as in (K) but for fresh frozen slices exposed to a 20 min, 1 h, or 4 h post-fix in 4% PFA at 4 °C or room temperature. (M) Schematic of a modified fresh frozen protocol that incorporates more aggressive post-fixation. (N-U) As in (B-I) but for mouse brain (N,O), lung (P,Q), duodenum (R,S), and skin (T,U) prepared using the protocol in (M). (V) The average counts per cell determined by MERFISH for the listed tissue for a fresh frozen protocol with either a standard or an enhanced post-fix. (W) The fraction of counts per cell associated with false positive controls (‘blanks’). Markers represent replicates while bars represent averages.

We prepared MERFISH samples for each of these four mouse tissues using this fresh frozen protocol, characterized gene expression with MERFISH, partitioned RNAs into cells, and explored the cellular diversity in these data (Figure 1B-I; Methods). We noticed a striking difference in the apparent quality of the data between these tissues. In brain and lung, major markers were found in the expected spatial locations (Figure 1B,D), RNA abundance was reproducible across replicates (Figure S2), and the expected cellular populations were largely resolved (Figure 1C,E). By contrast, duodenum and skin exhibited markedly reduced marker gene detection (Figure 1F,H), RNA abundance was less well reproduced across replicate measurements for the duodenum (Figure S2), and cell populations were poorly resolved in both (Figure 1G,I), suggesting that the measurement quality in these tissues was poor.

We next sought to quantify the differences in measurement performance between these tissues. As the targeted nature of image-based transcriptomics methods prohibits a meaningful comparison of metrics such as the average RNA counts per cell between different tissues, we instead quantified the ability of these measurements to define the cell populations expected within these tissues. To this end, we co-embedded these data with high-quality reference datasets^49–54^, transferred the reference labels to the MERFISH measurements, and used these reference-derived labels to explore the effective biological resolution of the different MERFISH datasets (Methods). Indeed, this analysis supported our qualitative observations: cell labels assigned via reference data clearly agreed with MERFISH clusters in the brain, to some degree in the lung, and relatively poorly in the duodenum and skin (Figure 1C,E,G,I); cell labels in the brain were assigned with greater confidence scores than in the other tissues (Figures S3, S4, S5, and S6; Methods); and the expected marker gene expression patterns for the transferred cell labels were better resolved in the brain and lung than in the duodenum and skin (Figures S3, S4, S5, S6). The biological resolution of cellular populations in the skin and duodenum was particularly poor with clear intermixing of functionally distinct cell type labels within different UMAP clusters and unclear expression of canonical cell type markers (Figure 1G,I; Figures S5, S6). Collectively, these results support the notion that there is a tissue-dependent difference in data quality for tissues prepared with fresh frozen protocols.

To explore the mechanisms that shape the observed tissue-dependent data quality, we turned to measures of the integrity of the RNAs within these samples, as we reasoned that tissue-dependent differences in the fragmentation of RNA might explain the differences in data quality. We harvested tissues rapidly, extracted RNA, and measured the RNA integrity number (RIN)—which is a measure of the degree of RNA fragmentation (Methods)^57^. We found that with proper harvest (Methods) all four tissues produced RIN values between 8 and 10, indicative of high RNA integrity (Figure 1J). As this observation suggested that substantial RNA fragmentation does not occur within these tissues during harvest, we next explored to what degree RNA degrades differentially during subsequent tissue processing. We performed RIN measurements on tissue sections obtained from cryoblocks for each of the four tissues. To simulate processing conditions, tissue slices were allowed to thaw for varying durations prior to RNA extraction and RIN assessment (Methods). Remarkably, we found that the RIN values decreased with time for all four tissues over the time scale of minutes (Figure 1J), suggesting active RNA degradation. However, there was clear disparity in RNA degradation rates, with RNA in the brain more stable than the lung which was in turn more stable than that of the duodenum and skin (Figure 1J). Indeed, for skin and duodenum the RIN dropped exceedingly quickly, plummeting to a value <5—indicative of highly degraded RNA—within 2 minutes.

As many RNases are compartmentalized within cells in order to regulate RNA degradation, we hypothesized that the freezing and slicing of tissues might damage membranes and other structures, releasing RNases which proceed to degrade RNA. To explore whether these tissue-dependent differences in RNA degradation rate represent tissue-dependent differences in the activity of endogenous RNases, we developed an on-slice RNase activity assay (Methods). With this assay, we observed RNase activity in cryosections of all four tissues (Figure 1K), with a relative hierarchy of activity consistent with the relative hierarchy of RNA degradation rates (Figure 1J) and of the apparent data quality (Figure 1C,E,G,I; Figures S2, S3, S4, S5, and S6). Collectively, these observations suggest that different tissues can degrade their own RNA remarkably quickly post sectioning and that tissue-dependent differences in RNase activity may explain the tissue-dependence of MERFISH performance for fresh frozen samples.

We next examined the degree to which the post-fixation step inactivates these endogenous RNases using the on-slice RNase activity assay. Surprisingly, we observed ample RNase activity in all four tissues after a 20-minute post-fixation at 4 °C (Figure 1L). Increasing the duration or the temperature of the post-fixation decreased RNase activity (Figure 1L), and 1 hour of room-temperature post-fixation was required to substantially reduce RNase activity across all tissues. As long post-fixation steps are uncommon for fresh frozen protocols, this observation suggests that many fresh frozen sample preparation protocols leave ample residual RNase activity in some tissues.

To determine if more complete inactivation of RNases could improve MERFISH performance in samples prepared with a fresh frozen protocol, we modified this protocol to include a 1-hour post-fixation at room temperature (Figure 1M) and explored MERFISH data quality in these four mouse tissues as above (Figure 1N-U; Figures S2, S3, S4, S5, and S6). Moreover, as these measurements target the same gene-tissue pairs as those characterized with a standard fresh frozen protocol, we could also compare basic performance metrics such as the average number of RNA molecules detected per cell (Figure 1V) and the fraction of false positive counts per cell (Figure 1W) between preparations within the same tissue. These measurements suggest that performance can be further improved in a tissue-dependent fashion with increased post-fixation. In the lung and skin, an increased post-fixation produced a noticeable increase in the RNAs detected per cell with a drop in the false positive rates (Figure 1V,W), and these improvements produced an improvement in the resolution of cell types (Figure 1Q,U; Figures S4, S6). By contrast, this extra fixation produced limited differences in performance in the brain and the duodenum (Figure 1O,S,V,W; Figures S3, S5) although likely for different reasons. The excellent cellular resolution in the brain with a light post-fixation suggests that there may be limited benefit to additional inactivation of the already low RNase activity. By contrast, the poor cellular resolution in the duodenum, independent of post-fixation conditions, may suggest that the high levels of endogenous RNase activity in this tissue may lead to such rapid RNA degradation that increasing the degree of fixation during the post-fixation is insufficient to rescue data quality: the RNA is already degraded by the time the PFA can sufficiently inactivate RNases.

### Improved MERFISH performance in fixed-frozen samples

Our measurements with a fresh-frozen tissue protocol suggest that endogenous RNase activity can be a major determinant of MERFISH data quality and that inactivation of these RNases can improve data quality in some contexts. If the loss of RNase compartmentalization in fresh frozen protocols leads to their active degradation of the sample, we reasoned that fixed-frozen protocols, in which the RNases would be inactivated prior to these steps, may offer improved performance. Thus, we next explored the effect of fixed frozen protocols on MERFISH data quality in these four mouse tissues.

In fixed frozen protocols, tissue blocks are rapidly dissected and then submerged in a fixative such as PFA for hours to inactivate and fix the tissue, the PFA is washed away, the tissue submerged in a cryoprotectant such as sucrose and then frozen, sectioned, and post-fixed as with fresh frozen samples (Figure 2A). The duration and temperature of the fixation step can vary widely^46–48,56^ with conditions generally optimized for the application and influenced by the relatively slow crosslinking kinetics of PFA in tissues^58^. Thus, we first sought to leverage our measurements of RNase activity to guide the fixation step towards more complete RNase inactivation. To this end, we leveraged a plate-based RNase activity assay (Methods), and we collected fixed frozen slices of the duodenum prepared with a PFA fixation step at 4 °C or room temperature for different durations. We selected the duodenum because it had both one of the highest levels of endogenous RNase activity of the tissues we profiled and its relatively thin structure would allow rapid penetration of PFA. Remarkably, we found substantial RNase activity even when the tissue was submerged in PFA for 3 hours with noticeably reduced but still detectable RNase activity even after fixation for 1 week at 4 °C or 48 hours at room temperature (Figure 2B,C). These observations suggest that aggressive fixation durations are required to substantially inactivate endogenous RNase activity in fixed frozen protocols.

**Figure 2.**
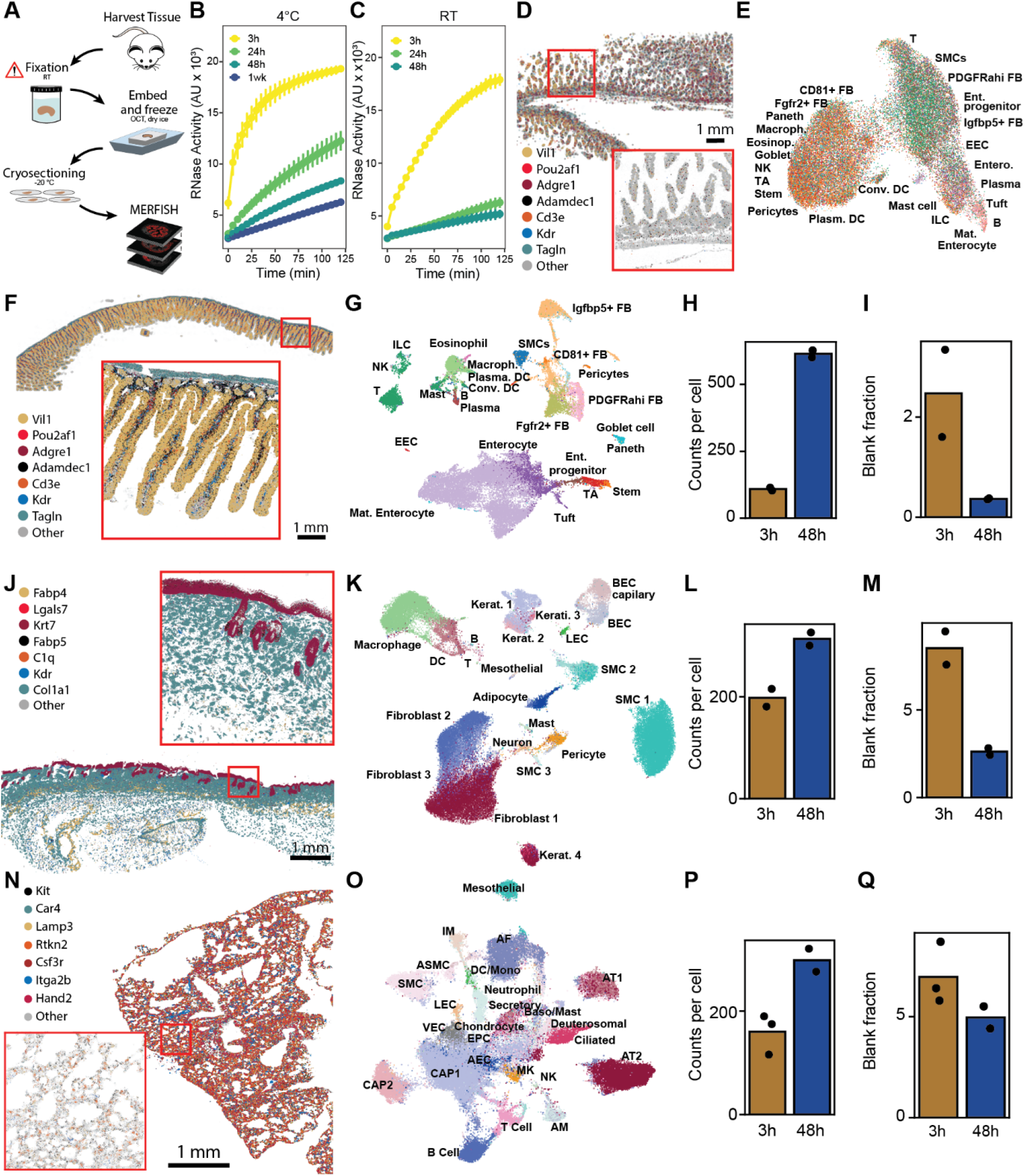
Extensive fixation inhibits RNase activity in fixed frozen protocols and improves MERFISH performance. (A) Schematic workflow for a fixed frozen tissue preparation for MERFISH. (B and C) RNase activity measured for a slice of mouse duodenum prepared as in (A) for the listed fixation durations prior to freezing for fixation at 4 °C (B) or room temperature (C). Error bars represent the standard error of the mean across three triplicate slices each from two separate tissue blocks. (D) Spatial distribution of 7 of 940 mRNAs measured with MERFISH in a longitudinal section of the mouse duodenum prepared as in (A) with a 3-hour, 4 °C fixation. (E) UMAP representation of the cellular diversity seen in the mouse duodenum with the protocol in (A) with a 3-hour, 4 °C fixation, colored by major cell type labels transferred to the data from a reference dataset (Methods). (F and G) As in (D and E) but for the mouse duodenum prepared as in (A) with a 48-hour, room-temperature fixation. (H and I) Average counts per cell (H) or average false positive fraction per cell (I) for the mouse duodenum prepared as in (A) but with a 3-hour, 4 °C fixation or a 48-hour, room-temperature fixation. Bars represent average over individual biological replicates represented as markers. (J-Q) As in (F-I) but for the mouse skin (J-M) or lung (N-Q) prepared as in (A) with a 3-hour, 4 °C fixation (L-Q) or a 48-hour, room-temperature fixation (J-Q).

To explore how reduced RNase activity would modulate the quality of MERFISH data, we performed MERFISH on fixed frozen blocks of the duodenum prepared with a 3-hour, 4 °C fixation (Figure 2D,E) and a 48-hour, room-temperature fixation (Figure 2F,G). As predicted given the high RNase activity observed after 3 hours of fixation at 4 °C, the quality of the MERFISH measurements from samples prepared with this fixation was poor and comparable to that observed for fresh frozen samples (Figure 1S, Figure 2E, Figure S5). Specifically, many labels associated with distinct cell types were intermixed within clusters in the UMAP (Figure 2E), these cell-type labels were applied with low confidence (Figure S5B), and canonical markers were not well expressed within these populations (Figure S5C).

By contrast, MERFISH measurements prepared with samples fixed for 48 hours at room temperature, where residual RNase activity was low (Figure 2C), were of decidedly higher quality (Figure 2F-I). While this improvement was reflected in a substantial increase in the number of RNA molecules detected per cell and a comparable decrease in the average false positive rate (Figure 2H,I), it was most strikingly seen in the improved resolution of cell types within these data. We observed clear coherence between the cell type labels and clusters on the UMAP (Figure 2G), these labels were applied with greater confidence (Figure S5B), and marker gene expression was clear (Figure S5C). Importantly, a wide variety of fine-grained cell types not clearly defined with previous preparation methods were now clearly defined, including not just major divisions within immune populations (e.g., macrophages, B cells, ILCs, T cells) but also specific immune subsets such as ILC1, ILC2, and ILC3, as well as CD4+ and CD8+ T cells (Figure 2G; Figure S5B,C).

Given this substantial performance increase observed with the duodenum, we next asked whether optimized fixed frozen protocols could improve the data quality in the other mouse tissues that we screened. Indeed, we found that for both skin and lung there was a clear improvement in data quality (Figure 2J-Q). As in the duodenum, basic performance metrics such as counts per cell and false positive levels were improved substantially with a 48-hour, room-temperature fixation relative to a 3-hour, 4 °C fixation for both skin and lung (Figure 2L,M,P,Q). Similarly, the confidence with which cell labels were assigned, the coherence with which these labels agreed with clusters in the UMAP, and the expression of marker genes were also markedly improved (Figure 2K,O; Figures S4, S6). By contrast, we observed only modest differences in the performance of MERFISH in the brain between samples prepared with either of the fresh frozen protocols or with fixed frozen protocols that employed a 3-hour, 4 °C or a 48-hour, room-temperature fixation (Figure 1C,O,V,W; Figure S7), again consistent with a model in which the low endogenous RNase activity of the brain is compatible with lightly fixed samples.

Collectively, these observations support the notion that tissue-dependent differences in endogenous RNase activity can have profound consequences for image-based transcriptomics data quality and that the differential inactivation of these RNases in fresh or fixed frozen sample preparation protocols can reveal these differences. Importantly, we show that measurements of the activity of RNases either through the speed with which RNA is degraded in slices of fresh frozen tissues (Figure 1J) or through measures of RNase activity within slices post fixation (Figure 1K,L; Figure 2B,C) can both stratify tissues and guide the optimization of tissue preservation protocols.

### Developing a PFA-compatible RNA preservation and tissue stabilization solution

While fixed frozen protocols with long PFA fixations were able to rescue data quality for the high endogenous RNase tissues we screened, we, nonetheless, were concerned that long incubations at room temperature may not be compatible with all tissues. For example, we and others have encountered some tissues or disease conditions for which RNA degrades rapidly and non-uniformly after tissue harvest^59–63^, likely far too quickly for the relatively slow inactivation of RNases by PFA. Moreover, the slow penetration of PFA into thicker tissue samples could allow residual metabolic activity within the tissue to remodel the transcriptome or degrade RNA, thereby introducing apparent cell states that are not an accurate representation of that observed in the original tissue. For these reasons, we sought to develop an RNA-aware histology approach that could rapidly stabilize the RNA within the sample and maintain this stabilization during long fixations. The rationale being that samples could be treated with a broad-spectrum RNase inhibitor that could rapidly penetrate and stabilize tissues. If this stabilizer were then compatible with chemical fixation, such as with PFA, the chemical inactivation could then be performed for the needed duration to fully inactivate RNases. We term this conceptual approach, Rapid Inhibition and Permanent Inactivation of Nucleases (RIPIN; Figure 3A).

**Figure 3.**
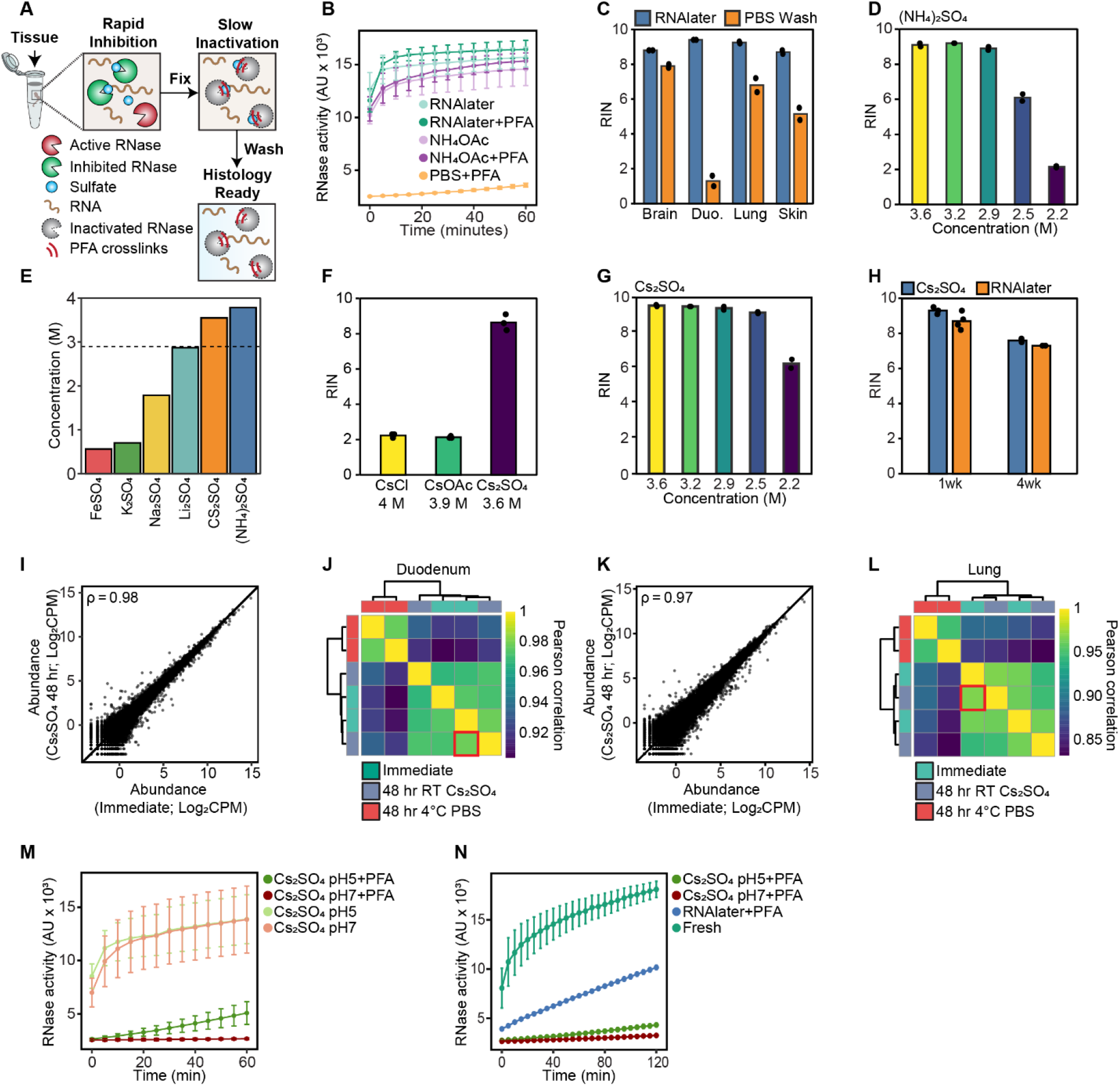
Developing a paraformaldehyde-compatible tissue preservation method to allow Rapid Inhibition and Permanent Inactivation of Nucleases (RIPIN). (A) Schematic representation of one implementation of the RIPIN framework, in which RNases are rapidly inhibited with concentrated sulfate solutions and then permanently inactivated with PFA modifications. (B) RNase activity determined by fluorescence in arbitrary units (AU) generated by a reporter in equal concentrations of RNase A treated or not treated with paraformaldehyde (PFA) in RNAlater, ammonium acetate (NH_4_OAc), or phosphate buffered saline (PBS). Markers and error bars represent the average and standard error across three triplicate measurements of two replicate treatments of RNase A. (C) RNA integrity number (RIN) measured for bulk RNA extracted from mouse brain, duodenum (Duo.), lung, or skin after storage in RNAlater for 48 hours at room temperature or after a 5 minute wash in PBS after this treatment. (D) RIN measured for bulk RNA extracted from mouse duodenum tissue blocks stored in an ammonium sulfate preservation solution at the listed concentration for 48 hours at room temperature. (E) Saturating concentrations of various sulfate salts in 20 mM EDTA at room temperature. The dashed line represents the minimum inhibitory concentration for ammonium sulfate. (F) RIN for RNA extracted from mouse duodenal tissue blocks stored in a preservation solution comprised of various cesium salts for 48 hours at room temperature. (G) RIN for RNA extracted from mouse duodenal tissue blocks stored in a cesium sulfate preservation buffer for 48 hours at room temperature with the listed cesium sulfate concentration. (H) RIN for RNA extracted from mouse duodenal tissue blocks stored in a cesium sulfate preservation buffer or RNAlater at room temperature for the listed times. (I) RNA abundance measured with bulk RNA sequencing in counts per million (CPM) for one murine duodenal tissue block stored in a cesium sulfate preservation buffer for 48 hours at room temperature versus that from a duodenal tissue block processed directly after harvest. ρ represents the Pearson correlation coefficient of the log_2_ CPM values. (J) Pairwise Pearson correlation coefficients for the log_2_ CPM values measured for two biological replicates each of duodenal tissue blocks harvested immediately or after 48 hours of storage in room temperature cesium sulfate preservation solution or 4 °C PBS. Samples are sorted via hierarchical clustering and colored by storage condition. The red box marks the comparison shown in (I). (K and L) As in (H) and (I) but for mouse lung tissue. (M) The fluorescence generated by a reporter of RNase activity for RNase A treated or not treated with PFA in cesium sulfate at the listed pH. Markers and error bars represent the average and standard error across three triplicate measurements of two replicate treatments of RNase A. (N) The fluorescence generated by a RNase activity reporter for slices of mouse duodenum processed via a fixed frozen protocol with fixation in a cesium sulfate preservation buffer at pH 5 or 7 or in RNAlater or prepared as fresh frozen samples (Fresh). Markers and error bars represent the average and standard error across three triplicate measurements of two tissue blocks.

To this end, we were inspired by the commercial RNAlater solution^34^, which has emerged as the gold standard for RNA preservation in tissue samples. RNAlater is primarily comprised of concentrated ammonium sulfate (∼3.6 M), which both rapidly penetrates tissues, due to its high osmotic strength, and inhibits enzymatic activity through non-specific adsorption of sulfate ions to protein^34,40^. Nonetheless, we did not anticipate RNAlater would be compatible with PFA fixation, as the ammonia in equilibrium with the ammonium ion is PFA reactive^64^. To explore this point, we developed an *in vitro* RNase A inactivation assay (Methods) to evaluate the ability to chemically inactivate RNases in different buffers. Using this assay, we found that while PFA was a potent inactivator of RNase A in phosphate buffered saline (PBS), it was ineffective in concentrated solutions of ammonium sulfate or ammonium acetate (Figure 3B). Moreover, we found that it is not possible to address these challenges by washing out the RNAlater, as many tissues showed rapid degradation of RNA after washout (Figure 3C). Nor was it possible to dilute the ammonium sulfate or RNAlater to a concentration at which PFA would be in molar excess of ammonium ions, as sulfate concentrations less than 2.5 M no longer protected RNA (Figure 3D; Figure S8A).

Ammonium sulfate is somewhat unique among the sulfate salts because of its very high saturating concentration (∼3.8 M, Figure 3E), which is the main reason why it was selected to be the active agent in RNAlater^34^. Of the readily available sulfate salts, most do not reach saturating concentrations high enough to expect substantial RNase inhibition (Figure 3D,E). The one exception is cesium sulfate^34^, which has a saturating concentration of ∼3.6 M (Figure 3E). As cesium is not expected to react with PFA, we proposed that cesium sulfate might be a suitable candidate for RIPIN.

Indeed, we found that an RNA preservation medium prepared with 3.6 M cesium sulfate reproduced the RNase inhibition and tissue stabilization features of RNAlater. First, we found that RNA extracted from mouse duodenum stored in a 3.6 M cesium sulfate preservation solution for 48 hours was of high quality, whereas RNA extracted from tissues stored in a similar preservation solution but with ∼4 M of cesium chloride or cesium acetate showed substantial degradation (Figure 3F), indicating that, like RNAlater, cesium sulfate can protect RNA and it is the sulfate that is responsible for RNase inhibition^34^. Second, we found that the inhibitory effects of cesium sulfate shared a similar concentration dependence as that observed for ammonium sulfate or diluted RNAlater (Figure 3D,G; Figure S8A). Third, we found that this cesium sulfate preservation buffer was capable of weeks-long preservation of mouse duodenal RNA at room temperature with identical performance to that observed with RNAlater (Figure 3H). Finally, to explore the ability of the cesium sulfate preservation buffer to preserve the transcriptional profile of tissues, we performed bulk RNA sequencing on mouse duodenal and lung tissue and compared the transcriptional profiles observed for tissues processed immediately or stored in the cesium sulfate preservation buffer at room temperature or PBS at 4 °C for 48 hours. For both tissues, we found that the transcriptional profiles measured for samples stored in cesium sulfate were essentially indistinguishable from that measured for samples extracted immediately after harvest (Figure 3I-L). Taken together, we conclude that cesium sulfate is a suitable substitute for ammonium sulfate for RNA preservation in tissue.

However, we did note one difference between the behavior of cesium sulfate and ammonium sulfate. It has been previously reported that the sample stabilization properties of ammonium sulfate require acidic pH^34^. While we reproduced this dependence for ammonium sulfate (Figure S8B), we found that the preservation capabilities of cesium sulfate did not have a strong pH dependence, and we found that neutral and even slightly basic solutions protected RNA (Figure S8C). This difference in performance is advantageous, as PFA fixation is less rapid at acidic pH^65^. Thus, the ability to work at neutral pH with cesium sulfate may enhance its compatibility with PFA.

To determine if this cesium sulfate preservation solution is PFA compatible, we first used the *in vitro* RNase A inactivation assay (Methods). We found that, unlike with RNAlater (Figure 3B), the activity of RNase A incubated with PFA in a cesium sulfate preservation buffer was substantially reduced relative to that not treated with PFA (Figure 3M). Moreover, these measurements confirmed that PFA-inactivation of RNase A was more potent in a cesium sulfate buffer at pH 7 relative to that of pH 5. To confirm that RNases could also be inactivated in tissue using the cesium sulfate preservation buffer, we prepared mouse duodenal tissue using a modified fixed frozen protocol (Figure 2A) in which the tissue was fixed with PFA for 48 hours in RNAlater or our cesium sulfate preservation buffer at pH 5 or pH 7. We then measured residual RNase activity within slices collected from these blocks. We found that samples fixed in cesium sulfate had a marked reduction in the endogenous RNase activity relative to those fixed in RNAlater (Figure 3N), confirming that our cesium sulfate preservation buffer was compatible with PFA fixation and might be a suitable approach for preserving tissues for spatial transcriptomic measurements via the RIPIN framework.

### RIPIN with a cesium sulfate preservation buffer produces high-quality MERFISH in all tissues

We next evaluated whether the cesium sulfate–based fixation method is compatible with MERFISH measurements. We prepared mouse brain, lung, duodenum, and skin using a modified form of the fixed frozen protocol (Figure 2A). Specifically, tissues were rapidly submerged into the cesium sulfate preservation buffer supplemented with PFA and fixed in this buffer for 48 hours at room temperature. With RNases inactivated via PFA fixation, we then could safely wash away the cesium sulfate, which would interfere with downstream histological steps. The samples were then prepared in sucrose, frozen, sliced, and stained for MERFISH as described above (Figure 2A; Methods).

Across all four tissues, we observed reproducible MERFISH measurements (Figure S2) with excellent data quality: marker genes were found in the expected locations; a high degree of cellular definition was seen in UMAPs; cell type labels were assigned with high confidence; and marker genes were clearly expressed (Figure 4; Figures S3, S4, S5, S6, S7). For all four tissues, we found that the average counts per cell were comparable to the highest of any of the other four preparation methods we considered (Figure 4D,I,N; Figure S7J) while the false positive levels were comparable to the lowest value observed with any of the other preparation methods (Figure 4E,J,O; Figure S7K).

**Figure 4.**
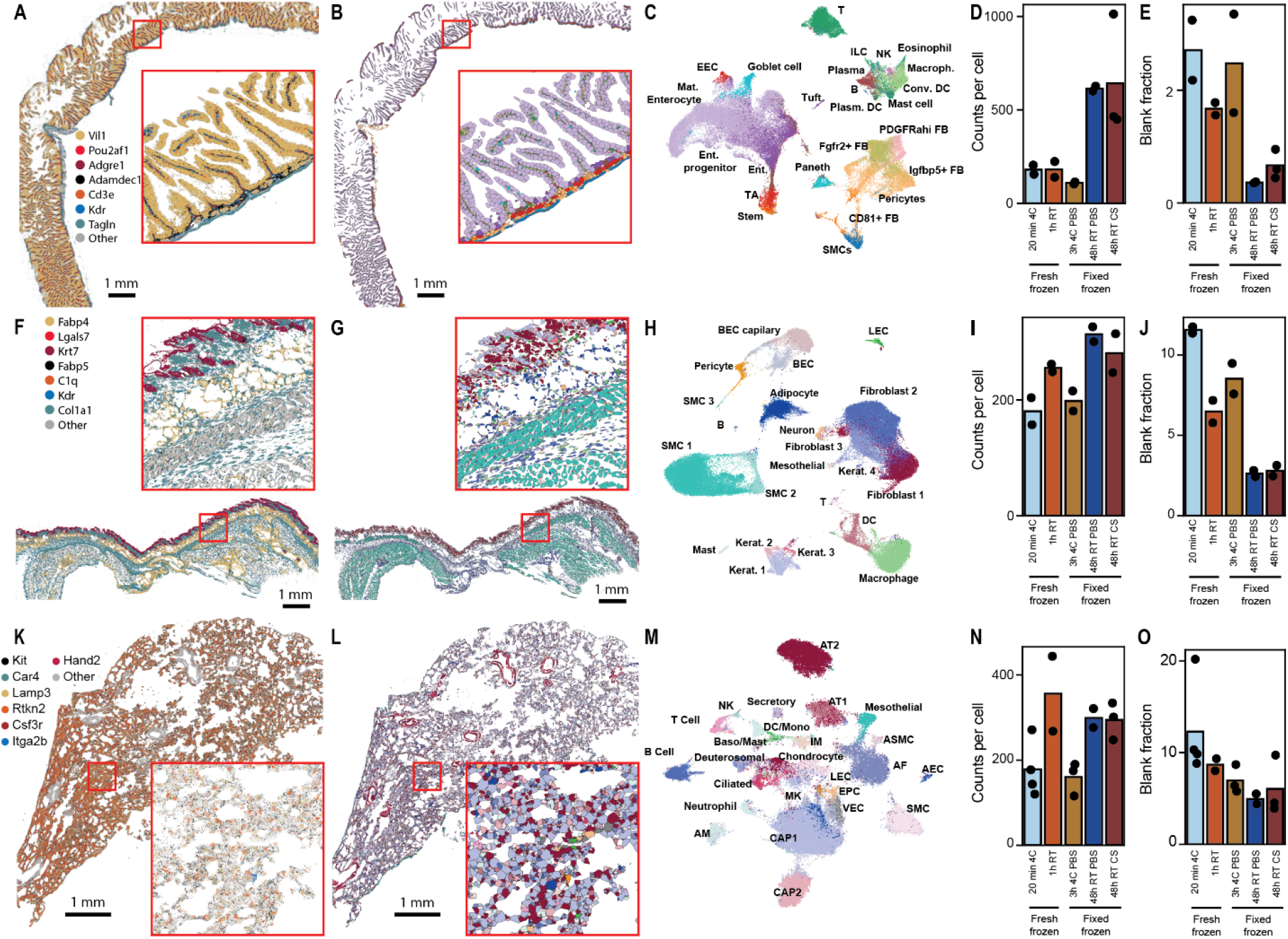
RIPIN with a cesium sulfate preservation buffer produces high-quality MERFISH in all profiled tissues. (A and B) Spatial distribution of 7 of 940 mRNAs (A) or of the identified cells (B) measured with MERFISH in a section of the mouse duodenum prepared with a fixed frozen protocol that leverages a 48-hour, room-temperature fixation in a cesium sulfate preservation buffer. (C) UMAP representation of the cellular diversity seen in two biological replicates of the measurements in (A), colored by major cell type labels transferred to the data from a reference dataset (Methods). (D and E) The average counts per cell (D) or false positive fraction per cell (E) for mouse duodenum samples prepared with each of the five listed protocols. Bars represent the average over biological replicates, which are listed as markers. Some values are reproduced from Figures 1 and 2 to facilitate comparison. (F-J) As in (A-E) but for the mouse skin. (K-O) As in (A-E) but for the mouse lung.

Most fixation methods produce some degree of change to the morphology of the tissue, and one would expect some such changes would arise from the high salt concentrations of the cesium sulfate preservation buffer. Although others have reported modest tissue morphology changes for some tissues soaked in RNAlater^66^, we, nonetheless, explored the modifications to tissue and cellular morphology in these samples. First, we noted that across all four tissues, the measured RNAs and cells were found in the expected locations and revealed an overall tissue morphology largely indistinguishable from that observed during fixation in PBS (Figure 4; Figure S7). To further explore potential morphological differences, we performed hematoxylin and eosin (H&E) staining on all four tissues fixed in a cesium sulfate preservation buffer or in PBS. High resolution imaging identified modest but noticeable differences that varied between tissues (Figure S9). Thus, as with all histology preparations, fixation in cesium sulfate does modify, to some degree, the native morphology of the tissue, and it will be important for researchers to both screen these defects in their tissue of interest as well as balance these defects over the facile preservation of RNA integrity.

As an important aside, the use of the same library-tissue combinations across multiple different preparation methods allowed us to quantitatively compare the effect of protocol choices on basic MERFISH performance metrics. These comparisons reinforced the central role that tissue-specific RNA degradation rates play in data quality as a function of preservation method. Specifically, we observed substantial increases in the average number of RNAs detected per cell and decreases in the fraction of these measurements attributed to false positives for the two tissues in which RNA degraded the most quickly: duodenum (Figure 4D,E) and skin (Figure 4I,J). Similar improvements were seen in the lung (Figure 4N,O), which showed a more intermediate degree of RNA degradation. By contrast, not only did increasing fixation not produce an improvement in the apparent data quality in the mouse brain—where RNA degradation rates were very low—we actually observed a modest but clear decrease in the number of RNAs detected per cell with increasing fixation (Figure S7J), suggesting, perhaps, that there is a cost to increasingly heavy fixation for tissues in which RNase activities are low enough to be neglected during tissue preparation.

### RIPIN with a cesium sulfate preservation buffer is compatible with clinical workflows

As heavy chemical fixation is a known confounder for the measurement of RIN or of the extraction of high-quality RNA for downstream analyses such as bulk RNA-sequencing, one additional advantage of RIPIN is the potential ability to temporally separate the stabilization of samples from their chemical fixation. In particular, we envisioned that these benefits could be particularly important for clinical workflows. Unlike tissues from model organisms, such as mouse, clinical samples may be far more unique, and it might be beneficial to collect matched bulk RNA-seq from portions of samples for which spatial transcriptomics would also be performed. Moreover, it may not be possible to always control the timing and nature of the tissue harvest in a clinical setting; thus, there may be instances in which RNA has degraded during harvest and before it can be stabilized in a preservation solution. In these cases, the ability to rapidly screen RNA integrity with RIN measurements could prove useful in selecting samples for spatial transcriptomics profiling.

To determine if RIPIN might offer these advantages, we explored the ability to delay the addition of PFA without comprising the performance of MERFISH measurements (Figure 5A). To this end, we collected mouse duodenum and immediately submerged it in the cesium sulfate preservation buffer. We held this tissue at room temperature for 24 hours, dissected a portion for RIN characterization, and then added PFA to fix the remaining portion for 48 hours. This residual tissue block was then prepared with the fixed frozen protocol (Figure 2A). We characterized cryosections of this block with MERFISH and observed that the measured cells co-integrated with those measured from samples immediately fixed in the cesium sulfate preservation buffer (Figure 5B) and, as expected, demonstrated the same high cellular resolution (Figure 4C and Figure 5C) as well as the same counts per cell as those samples (Figure 5D). As expected, we found that the RIN measured after 24 hours of stabilization and prior to PFA fixation matched that of tissue processed immediately (Figure 5E). Thus, RIPIN allows samples to be stabilized, tissue subsets to be collected and processed for PFA-sensitive RNA applications, and then fixed afterwards, with no obvious detriment to the downstream spatial transcriptomics data quality.

**Figure 5.**
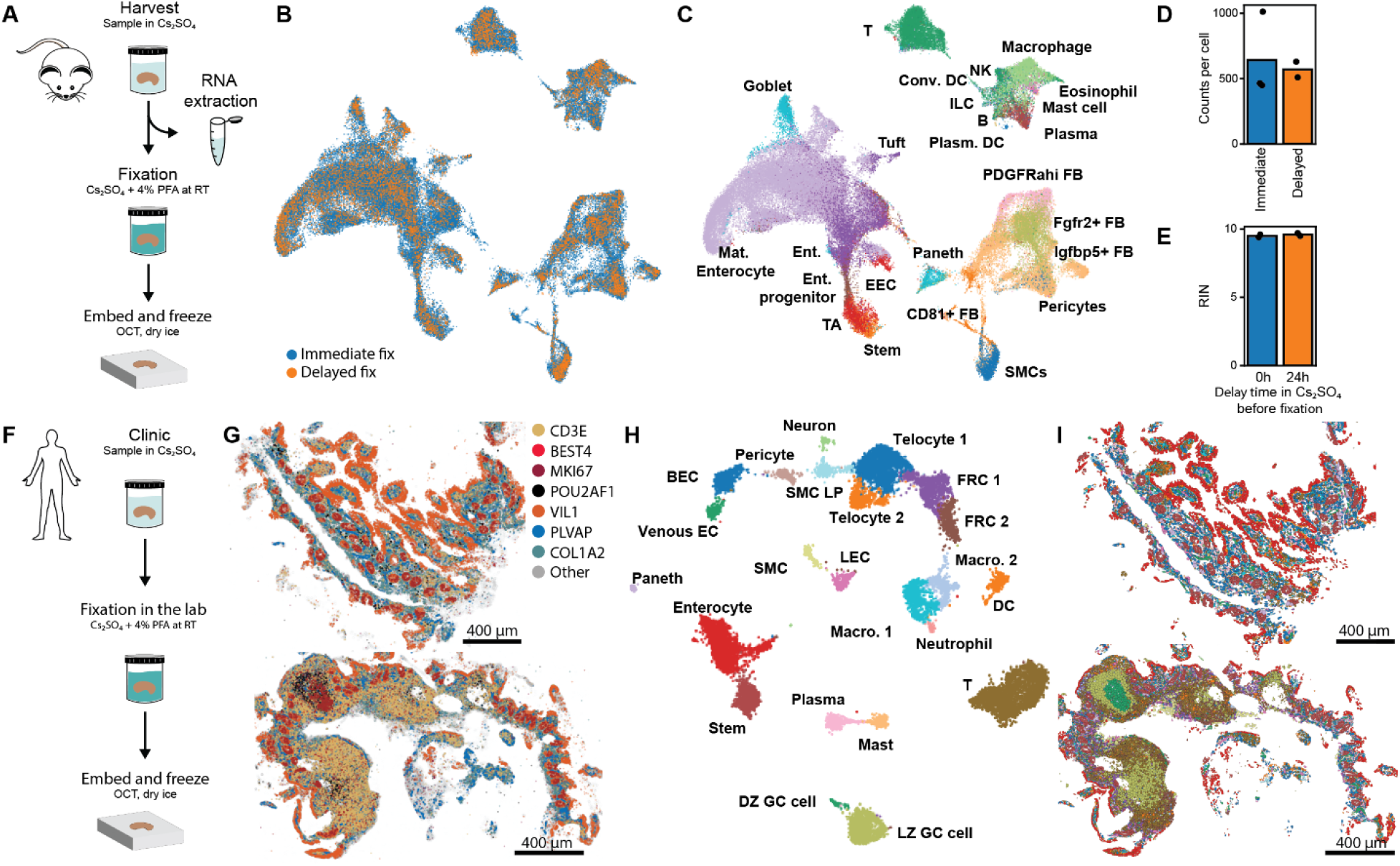
RIPIN is compatible with clinical workflows. (A) Schematic workflow for cesium sulfate–based tissue collection, decoupling RNA preservation from fixation and enabling PFA-sensitive RNA quality measurements within the same sample. (B and C) UMAP representation of the cellular diversity seen in the mouse duodenum fixed for 48 hours at room temperature in a cesium sulfate preservation buffer with either the immediate addition of PFA or after 24 hours of room temperature storage in a cesium sulfate preservation buffer without PFA, colored by preparation method (B) or cell type label (C). (D) The average counts per cell for mouse duodenum samples prepared with a 48-hour, room temperature fixation in the cesium sulfate preservation buffer with or without an initial 24-hour incubation in cesium sulfate preservation buffer. Values labeled as *Immediate* are reproduced from Figure 4D to facilitate comparison. (E) RIN for mouse duodenal RNA extracted immediately or from samples stored for 24 hours at room temperature in the cesium sulfate preservation buffer prior to PFA addition. (F) Schematic workflow of clinical samples collection in a cesium sulfate preservation buffer, where samples are immediately stabilized in the procedure room and then fixed sometime later through the addition of PFA. (G) Spatial distribution of 7 of 737 mRNAs measured with MERFISH in ileal biopsies from two Crohn’s disease patients. (H) UMAP representation of the cellular diversity seen in the measurements in (G) colored by cell type label. (I) Spatial distribution of the cells seen in the biopsies in (G) colored as in (H).

To demonstrate that this approach can be integrated into clinical workflows, we applied it to the collection of human intestinal biopsies from Crohn’s disease patients. During a standard diagnostic endoscopy procedure, pinch biopsies were collected, submerged into cesium sulfate preservation buffer—with no added PFA—and incubated over the ∼15-45-minute duration of the procedure. These samples were then brought to a research laboratory environment, where PFA was added roughly 1-3 hours after the samples were collected (Figure 5F). The samples were then fixed for 48 hours at room temperature and prepared using the fixed frozen protocol (Figure 2A). In parallel, we developed a ∼737-gene human gut-focused MERFISH library and characterized sections of biopsies collected in this fashion with this library. Supporting the efficacy of the RIPIN approach in this clinical setting, the quality of the MERFISH data were high. We observed dense expression of marker genes in the expected locations, excellent cellular diversity within UMAP representations of the data, and cell type distributions and tissue morphologies expected for intestinal biopsies (Figure 5G-I). These results confirm the ability to separate the stabilization and fixation of the sample and to integrate RIPIN into clinical workflows. As cesium sulfate, unlike PFA, is considered non-toxic and non-hazardous, the ability to remove a chemical of concern from the clinical environment may represent another practical benefit.

### Cesium sulfate–based fixation is compatible with paraffin embedding

Finally, we investigated whether cesium sulfate–based fixation is compatible with paraffin embedding, one of the most widely used tissue embedding methods. Sectioning paraffin embedded samples offers key advantages over cryosectioning, as the rigid paraffin matrix facilitates sectioning of even soft or fatty tissues without the chatter, curling, or tearing that can occur in challenging tissues during cryosection. Moreover, paraffin sectioning does not require cryogenic temperatures and is routinely performed at room temperature. In formalin fixed paraffin embedding (FFPE) protocols, samples are fixed in formalin for various durations, dehydrated in a graded ethanol series, and then embedded in paraffin wax (Figure 6A). Sections are prepared with a microtome, placed on coverslips, deparaffinized by melting the wax and solubilizing it with organics such as xylene, and then rehydrated in a graded ethanol series. From there, standard MERFISH protocols can be applied (Figure 6A).

**Figure 6.**
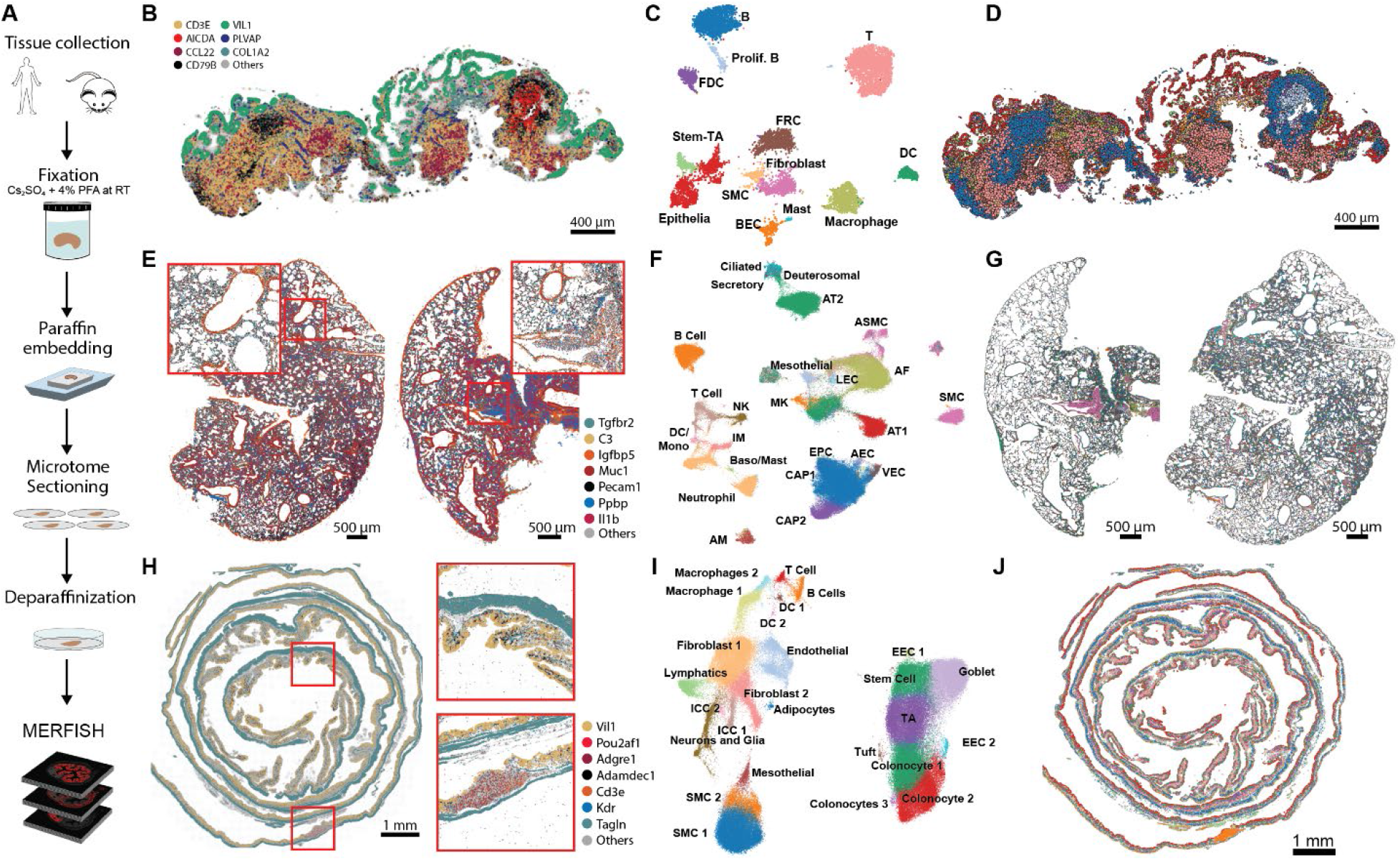
RIPIN is compatible with paraffin embedding. (A) Schematic for a paraffin embedding workflow for samples fixed in the cesium sulfate preservation buffer (Methods). (B) Spatial distribution of 7 of 737 mRNAs measured with MERFISH in an ileal biopsy from a Crohn’s disease patient prepared as in (A). (C) UMAP representation of the cellular diversity seen in the biopsy. (D) Spatial distribution of the cells in (B) colored as in (C). (E-J) As in (B-D) but for the mouse lung measured with the 940-gene library (E-G) or the mouse colon measured with the 940-gene [Cell 2024] library (H-J).

As the paraffinization and deparaffinization processes involve steps that might naturally inhibit or inactivate RNases—namely the dehydration in alcohol and the organic extraction of melted paraffin—we first asked whether these steps might have the added benefit of removing RNase activity. To first test alcohol treatment, we started with a common alcohol-based fixative solution. We harvested mouse ileum and incubated it overnight in the alcohol-based fixative PAXgene^67^, and then extracted RNA immediately or after a wash and incubation in PBS. RIN measured while the tissue remained in the preservative solution was near optimal, whereas RIN values obtained after the tissue was washed with PBS for an hour were markedly reduced (Figure S10A). These findings suggest that alcohol treatments do not permanently inactivate RNases, likely because many RNases readily refold upon re-exposure to aqueous conditions.

We then tested whether the entire paraffin embedding and de-embedding process might inactivate RNases and contribute to RNA preservation, as these processes involve organic solvents. Mouse duodenal tissues were harvested, lightly fixed for 3 hours at 4 °C to preserve RNase activity or heavily fixed for 48 hours at room temperature to inhibit RNase activity, and prepared in paraffin (Figure 6A; Methods). After slicing and deparaffinization, we observed ample RNase activity on slices that were lightly fixed relative to those that were heavily fixed (Figure S10B-D), indicating that the process of paraffinizing and deparaffinizing offers no direct benefit for RNase inactivation.

This observation also raises an important concern for the use of pre-existing tissue microarrays (TMA) or other archival paraffin embedded samples for spatial transcriptomics measurements, as it is possible that such samples were not fixed sufficiently to fully inactivate RNases prior to paraffin embedding. To explore this possibility, we measured the RNase activity of a series of mouse tissues from commercial TMAs prepared from FFPE samples with our previously established RNase activity assay. Remarkably, we detected reproducible and substantial residual RNase activity in a subset of tissues within these TMAs (Figure S10E). This observation suggests that screening such samples with the simple on-slice RNase activity assay could help guide the selection of samples for spatial transcriptomic characterization.

We next prepared a diversity of mouse and human tissues using a cesium-sulfate-preserved, formaldehyde-fixed, paraffin-embedded protocol (Figure 6A) to determine whether we could obtain high-quality MERFISH measurements with these preparations. Specifically, we characterized human intestinal biopsies (Figure 6B-D), mouse lung (Figure 6E-G), whole Swiss rolls of the mouse colon (Figure 6H-J), and coronal sections of the mouse brain (Figure S7L-N). In all cases, we observed clear spatial distributions of established marker genes and excellent transcriptional and spatial definitions of the expected cell populations. Importantly, we noticed that paraffin embedding offered an additional advantage for processing slices of heavily fixed samples. Some tissues displayed increasingly poor adherence to coverslips with heavy fixation in fixed frozen protocols, and these adherence issues could generate tissue defects. Paraffin embedding substantially improved coverslip adherence and reduced or eliminated these defects. Thus, we conclude that RIPIN as implemented with our cesium sulfate preservation buffer is readily incorporated into paraffin embedding workflows.

## DISCUSSION

Spatial transcriptomics technologies promise to integrate the strengths of histology and genomics to power biological discovery, but they also inherit the sample preparation challenges associated with both. The development of these techniques and their early success in fresh frozen mouse brain^11,19,20,22,23,68,69^ may have led to an underappreciation of the challenges of sample preparation. As these methods are now being widely adopted across a greater diversity of tissues, disparities in data quality between tissues are being discovered^27–30^. Nonetheless, the sources of this variation remain poorly understood, and assays to guide optimization of tissue preparation across diverse tissue types are still lacking. In this study, we address these challenges by systematically evaluating how commonly used histology preparation methods influence RNA stability and, in turn, image-based spatial transcriptomics data quality across a diverse range of tissues. Our findings demonstrate that endogenous RNases play a significant role in shaping data quality by driving RNA degradation during key stages of sample processing, in a tissue-dependent manner. Notably, we show that high-quality MERFISH measurements are closely associated with effective RNase inactivation, which is typically achieved through extensive fixation. Furthermore, we introduce simple assays to test for endogenous RNase activity, which can be used to optimize fixation conditions for high-quality spatial transcriptomic measurement in a wider range of tissues than tested in this work.

While we have focused on one image-based method here, we expect that residual endogenous RNase activity will impact data quality for all image- and capture-based spatial transcriptomics technologies and that optimizing tissue preservation methods to reduce this activity at critical stages of sample preparation may enhance data quality across all platforms. To this end, our work suggests that new tissues should be optimized with a multi-step process to guide sample preparation for high-quality spatial transcriptomic measurement. Where possible, we recommend preparing fresh frozen tissue blocks and measuring RIN versus time for slices taken from these blocks. If the RNA is largely stable over long durations, as we observed for the mouse brain, then fresh frozen protocols may be the best options for that tissue. By contrast, if RNA degrades quickly, then heavy fixation of the tissue block is likely to produce slices with more intact RNA. In this context, we further recommend measuring the residual RNase activity within individual slices and optimizing fixation duration to minimize this activity.

While all spatial transcriptomic methods will be sensitive to RNA fragmentation, it is important to acknowledge that different methods may have different sensitivity to RNA integrity and, thus, the benefit scale associated with more complete RNase inactivation seen with MERFISH may not be shared with other methods. Indeed, we would anticipate that image-based methods that use a small number of probes per RNA, i.e. Xenium^13,14^, STARmap^11,12^, or CosMx^24^, may be able to tolerate a greater degree of RNA fragmentation than those that use multiple tens of probes per RNA, i.e. MERFISH^17–21^ and seqFISH^22,23^. Similarly, spatial capture methods, which often require sequencing only short fragments from the 3’ end of RNAs, may also be less sensitive to RNA fragmentation. Indeed, a platform-specific sensitivity to RNA fragmentation coupled with residual RNase activities that are both tissue and preparation dependent may explain the inconsistent performance differences in image-based platforms observed across a range of recent benchmarking studies^27–30^. Importantly, our work underscores the challenge of drawing general conclusions on the relative performance of different image-based transcriptomic chemistries from benchmarking studies that do not control for the substantial differences in RNA degradation that we observe here for different tissues.

In parallel, the modern histology preparation toolbox is remarkably varied, comprising a diversity of fixation buffers and methods in combination with diverse embedding and sectioning methods. This diversity has largely arisen to solve a wide variety of tissue-specific challenges. However, our work now reveals a critical underexplored element in this toolbox: RNA-aware histology methods. The lack of these methods is perhaps not surprising, as many of the techniques in wide use were developed long before the current era of genomics. In this context, we anticipate that our RIPIN framework—rapidly stabilize the RNAs within samples and then permanently inactivate the RNases—may represent an important concept in the development of RNA-aware histology methods. As an initial contribution, we introduced here a cesium sulfate preservation buffer that stabilizes RNA within samples in a manner comparable to the widely used RNAlater solution. Notably, we show that this buffer can support PFA fixation and can, thus, be used to rapidly stabilize RNAs and maintain the inhibition of RNase activity during the long PFA fixations we find are necessary to inhibit RNases in many tissues. We have found that the cesium sulfate preservation buffer offers many advantages, including high-quality MERFISH in all profiled tissues, the ability to stabilize RNAs and characterize samples with fixation-sensitive RNA methods prior to fixation, and broad compatibility with tissue embedding and sectioning methods. More broadly, we have found that the use of the cesium sulfate preservation buffer is more robust for less experienced researchers, generating high-quality tissue measurements more rapidly for those with limited tissue harvest and preservation experience.

Nonetheless, there are several disadvantages to the cesium sulfate preservation solution. First and foremost, fixation in a high salt solution can introduce morphological artifacts, and, while these are modest for the tissues we profiled here, they could be more severe for other tissues. Notably, other preservation methods, such as fixation in PBS-PFA, are also known to introduce tissue-specific morphological differences. Thus, such histological defects are not unique to the cesium sulfate preservation buffer. Moreover, the vast majority of current spatial transcriptomic studies currently make limited use of the fine features of cellular morphology; thus, we anticipate that the benefits of robust production of intact RNA during histological preparation may offset concerns with these modest morphological defects for many studies. However, as this field begins to better integrate measures of tissue and cellular morphology with transcriptomic measurements, this issue may be increasingly important. In this context, alternative approaches to the RIPIN framework could be interesting to explore. For example, while we demonstrated that alcohol-based fixatives are insufficient for permanently inhibiting RNases, they are potent inhibitors of RNases and are commonly used in contexts where specific morphological features in a tissue must be preserved. Thus, it may be possible to combine these solutions with chemical fixatives to irreversibly inactivate RNases, implementing an alternative RIPIN formulation that may be better suited for morphological preservation in some situations. Indeed, the modern histology toolbox is filled with many variant fixation solutions designed specifically to preserve some aspect of the morphology of individual tissues. We envision that RIPIN may provide a framework to modify or augment these methods to both obtain optimal morphology and RNA integrity in specific tissues.

More broadly, our work clearly illustrates the degree to which spatial transcriptomic data quality—and thus the types of biological conclusions one can draw—depends critically on the RNA integrity and, therefore, tissue preparation methods used. By drawing attention to this critical point, we anticipate that our work may accelerate the development of optimized RNA-aware protocols for a variety of tissues and that these protocols will provide the spatial transcriptomics data quality that is necessary to reveal previously unknown aspects of cell and tissue biology in a wide variety of contexts.

## Lead contact

Requests for further information and resources should be directed to and will be fulfilled by the lead contact, Jeffrey Moffitt.

## ACKNOWLEDGEMENTS

We thank members of the Moffitt and Cantor laboratories for helpful discussions. We thank N. Twumasi-Ankrah for help with illustrations. Portions of this research were conducted on the O2 High Performance Compute Cluster, supported by the Research Computing Group, at Harvard Medical School. We thank the Harvard Cancer Consortium in Boston, MA, for the use of the Rodent Histopathology Core, which provided paraffin embedding, microtome sectioning, and H&E staining services in addition to general histology advice and guidance. The Harvard Cancer Consortium is supported in part by a NCI Cancer Center Support Grant # NIH 5 P30 CA06516. We are also deeply grateful to the anonymized donors of human tissue. This study was supported by grants from the Leona M. and Harry B. Helmsley Charitable Trust (to J.R.M. and S.B.S.), the National Institutes of Health (R01GM143277 to J.R.M.; R01HL159106 to J.R.M. and A.B.C.; R01NS113890 to S.S.; P30DK034854 to S.B.S.; and 5T32HL007574-36 which supported C.A.R.-L.), from The G. Harold and Leila Y. Mathers Charitable Foundation (to J.R.M. and A.B.C.), from the Veterans Administration (VA Merit Award I01 BX006131 to S.S.), and from the Charles A. King Postdoctoral Research Fellowship Program from the Bank of America Co-trustees (to P.C.).

## AUTHOR CONTRIBUTIONS

P.C., C.A.R-L., H.Z., and J.R.M. conceived the study and designed experiments. P.C., C.A.R-L., H.Z., and K.K. performed experiments. P.C., C.A.R-L., H.Z., and B.R.W. developed reagents. P.C., C.A.R-L., and H.Z. analyzed data. P.C., C.A.R-L., and S.B.S obtained intestinal biopsies. S.S., S.B.S., A.B.C., and J.R.M. supervised work. P.C., C.A.R-L., H.Z., and J.R.M wrote the manuscript. P.C., C.A.R-L., H.Z., K.K., B.R.W., S.S., S.B.S., A.B.C, and J.R.M revised the manuscript.

## DECLARATION OF GENERATIVE AI AND AI-ASSISTED TECHNOLOGIES IN THE WRITING PROCESS

During the preparation of this work, the authors used Anthropic Claude-Sonnet-4.6 in order to identify typos and formatting issues. After using this tool, the authors reviewed and edited the content as needed and take full responsibility for the content of the publication.

## METHOD DETAILS

### Animals

C57BL/6J mice (JAX000664) were purchased from Jackson Labs and used at 10-15 weeks of age. Mice were housed in the Harvard Center for Comparative Medicine (HCCM) facility before euthanasia and tissue harvest. All mouse experiments were performed in compliance with NIH guidelines and were reviewed and approved by the Harvard Institutional Animal Care and Use Committee (IACUC) under protocol IS00003215.

Mice were euthanized with either isoflurane (Patterson, 07-890-8115) overdose or CO_2_ asphyxiation followed by cervical dislocation. Tissue was rapidly dissected and then prepared according to one of the protocols described below.

### Mouse tissue harvest

To profile the tissue dependence of MERFISH quality for different tissue preparation and preservation methods in a reproducible fashion, we selected four mouse tissues for characterization: brain, lung, duodenum, and skin. In addition, we profiled a range of different histological tissue preservation and preparation methods: a fresh frozen protocol, a cold fixed frozen protocol, a room temperature fixed frozen protocol, and a room temperature fixed frozen protocol that leverages a cesium sulfate preservation solution. Minor modifications were made to tissue harvest based on the preservation method.

To harvest the brain, the skull of euthanized mice was dissected, and the entire brain was quickly removed and placed in a preservation solution. For the fresh frozen and cold fixed frozen protocols, this buffer was ice-cold 1× Phosphate Buffered Saline (PBS; Thermo, AM9625) or an ice-cold PBS-based fixation solution. For the room temperature fixed frozen protocol, this buffer was a room temperature cesium-based preservation and fixation solution. The fixation and preservation solutions are described below.

To harvest the lung, the rib cage was opened at the sternum, lungs were removed, and the trachea severed. Lungs were then briefly perfused with either ice-cold 1× PBS (for the fresh frozen protocol or the cold fixed-frozen protocol) or room temperature 1× PBS (for all room temperature fixation protocols). This perfusion was necessary to both improve RNA quality and to provide better sectioning quality.

To harvest the duodenum, the abdominal cavity was opened, and the duodenum was injected with 2 mL of 1× PBS or the cesium sulfate preservation solution with a 27G needle (Becton Dickinson, 305109). The exterior of the duodenum was washed with 1 mL of this same solution, and then the duodenum was dissected and placed directly into ice-cold 1× PBS (for fresh frozen preparation) or the fixation solution used for the fixed frozen protocol at the temperature at which the fixation was conducted. The flushing of the duodenum was necessary to protect the tissue from pancreatic fluids, which are rich in RNases.

To harvest the skin, the fur on the abdomen of the mouse was first removed using an electric trimmer. The skin was then briefly coated with 70% ethanol. We found that both the removal of the fur and the ethanol treatment were necessary to increase the wettability of the tissue and to obtain rapid contact with buffers. The skin was then cut from the abdomen and placed in ice-cold 1× PBS (for fresh frozen) or the fixation buffer at the temperature of the fixation (for all other protocols).

We also leveraged mouse colon to demonstrate the ability of the cesium sulfate preservation buffer to be combined with paraffin embedding. To harvest the colon, the abdominal cavity was opened, and the entire colon was carefully dissected. The colon was then incised longitudinally, rapidly rinsed with a cesium sulfate preservation buffer and subsequently fixed in the same solution used for the fixed-frozen protocol at room temperature. Following fixation, the colon was rolled into a Swiss roll configuration for further processing.

### Cesium sulfate and ammonium sulfate preservation buffers

The cesium sulfate preservation buffer was prepared by dissolving 181 g of cesium sulfate (Chem-Impex, 10294-54-9; Thermo, 218702500) in 96 mL of water. This solution was supplemented with 4 mL of 0.5 M EDTA (Thermo, AM9262), and the final pH of the solution was adjusted to 7.0, unless otherwise specified, through the addition of either 1 M H_2_SO_4_ (Sigma, 258105) or with 10 N NaOH (Sigma, 72068) for initial adjustment and 1 N NaOH for fine tuning. The final volume of this solution was ∼140 mL, bringing the final concentration of cesium sulfate to ∼3.6 M. The cesium sulfate preservation buffer could be stored at room temperature for at least 3 months without obvious loss in performance. When this buffer was supplemented with paraformaldehyde (PFA) to fix samples, PFA was brought to its final concentration of 4% (w/v) by dilution of 32% (w/v) PFA (Electron Microscopy Sciences, 15714), which produced a corresponding dilution of the cesium sulfate preservation buffer. All water used in this work was deionized, RNase-free and DNase-free water prepared by MilliQ water purification system (Millipore, Synergy) with a biopack polisher (Sigma, CDUFBI001).

The active ingredient of the commercial RNA preservation solution, RNAlater (Invitrogen, AM7021), is ammonium sulfate^34^. While the manufacturer does not disclose the full composition of RNAlater, we created an ammonium sulfate preservation solution that aims to emulate its composition based on published values^34^. Briefly, 47 g of ammonium sulfate (Sigma-Aldrich, A2939-1KG) was dissolved in 60 mL of 20 mM EDTA (prepared from 0.5 M EDTA stock) and stirred at room temperature for 2 hours. The solution was then brought to a final volume of 100 mL with 20 mM EDTA, which corresponds to a final concentration of ammonium sulfate of ∼3.6 M. The pH was then adjusted to 5.0 using 1 M H_2_SO_4_, unless otherwise specified. When fixation was needed, PFA was added from a 32% (w/v) stock solution to achieve a final concentration of 4%. Where listed as RNAlater, we used commercial RNAlater diluted in water or pH adjusted with 10 N NaOH as appropriate. Where listed as ammonium sulfate, we used this home-built ammonium sulfate solution.

For control experiments, we created versions of the preservation solution composed of cesium acetate, cesium chloride, or ammonium acetate. Because sulfate is the active component associated with RNase inhibition, these solutions are not expected to preserve RNA. Cesium acetate (Sigma, 450154-25G) was prepared by dissolving 4.8 g in 5 mL of 20 mM EDTA, followed by pH adjustment to 7.0 with 1 M H_2_SO_4_. The final volume was ∼6.4 mL, yielding a concentration of ∼3.9 M. Cesium chloride (Sigma, 289329-100G) was prepared similarly by dissolving 4.2 g in 5 mL of 20 mM EDTA, and then adjusting the pH to 7.0 with 1 M H_2_SO_4_, resulting in a final volume of 6.2 mL and a concentration of ∼4.0 M. Ammonium acetate (Sigma-Aldrich, A1542-250G) was prepared by dissolving 5.6 g of salt in a final volume of 10 mL of 20 mM EDTA to achieve a concentration of ∼7.2 M, corresponding to an equivalent ammonium ion concentration as ∼3.6 M ammonium sulfate. The pH was then adjusted to 7.0 using 5 M acetic acid (prepared from glacial acetic acid [Fisher, A38-500]). When samples were fixed in these solutions, a final concentration of 4% (w/v) PFA was created by diluting the 32% (w/v) stock solution directly into these buffers as described above.

To determine the solubility of additional potential sources of sulfate, we incrementally added each sulfate salt to 10 mL of 20 mM EDTA until no further dissolution was observed. Additional 20 mM EDTA was then added to fully dissolve the remaining solid. The final solution volume was measured and used to calculate the saturated concentration for each salt. Using this approach, we found that 2.5 g of iron(II) sulfate heptahydrate (Sigma, F7002) dissolved in a final volume of ∼16 mL; 4.1 g of lithium sulfate (Thermo, 447552500) in ∼13 mL; 3.8 g of sodium sulfate (Sigma, 239313) in ∼15 mL; 2.2 g of potassium sulfate (Sigma, P0772) in ∼18 mL; 18.1 g of cesium sulfate in ∼14.1 mL; and 5 g of ammonium sulfate in ∼10 mL. These values reflect the relative solubility of each salt under the tested conditions and may vary depending on factors such as salt lot, solution composition, and pH.

### Fresh- or fixed-frozen tissue block preparation

For the fresh frozen protocol, tissue blocks were created by placing the tissue in an ice-cold plastic cryomold (Fisher, 22-363-554) filled with an initial layer of optical cutting temperature media (OCT; Fisher, 1437365). The mold was then filled, covering the tissue completely, with OCT. The sample was then snap-frozen on dry ice.

For the fixed frozen protocols, samples were incubated in either 1× PBS or the cesium sulfate preservation solution supplemented with 4% (w/v) PFA. For cold fixation, samples were incubated for 3 hours at 4 °C. For room temperature fixation, samples were incubated in the fixation solution for 48 hours at room temperature. In both cases, the samples were incubated in a 50 mL falcon tube with at least 10-fold greater volume of the fixation solution than the tissue volume, and the samples were gently rocked to ensure solution mixing for the duration of the fixation. Samples prepared in 1× PBS were then directly transferred to a solution of 14 mL of 30% (w/v) sucrose (Thermo, 419760010) prepared in water (duodenum and skin) or 1× PBS (brain and lung), gently rocked at 4 °C for ∼24 hours. Samples prepared in the cesium sulfate preservation solution were washed five times for 10 minutes each in 50 mL of water (duodenum, skin, and brain) or 1× PBS (lung) to remove the excess cesium sulfate before transferring to the 30% (w/v) sucrose solution prepared in water (duodenum, skin, and brain) or 1× PBS (lung). Samples were incubated in the sucrose solution, with gentle rocking, for ∼24 hours at 4 °C. They were then transferred to a cryomold and embedded in OCT, as described above.

### Human tissue collection and processing

Biopsy samples from the terminal ileum and colon were obtained from pediatric patients with Crohn’s disease (CD) at the time of diagnostic endoscopy. The diagnosis of CD had been previously established by the treating clinicians based on a combination of endoscopic, histopathologic, and clinical criteria. Written informed consent was obtained from all participants in accordance with the guidelines of the Boston Children’s Hospital Institutional Review Board (Protocol P00000529). Biopsies were immediately placed at room temperature into 2 mL tubes containing 2 mL of the cesium sulfate preservation solution. Samples were transported to the laboratory within 30 to 60 minutes and subsequently transferred into fresh 2 mL tubes containing 2 mL of cesium sulfate preservation solution supplemented with 32% (w/v) PFA to a final concentration of 4% (w/v). Samples were fixed in this solution for 48 hours at room temperature. Following fixation, samples were washed five times for 10 minutes each in 50 mL of deionized water. Tissues were then either processed for paraffin embedding as described below or cryoprotected in 30% (w/v) sucrose in deionized water overnight at 4 °C and subsequently embedded in OCT as described above.

### Fresh frozen sample preparation for MERFISH

Coverslips (Bioptechs, 40-1313-03193) were silanized and coated with poly-d-lysine (Gibco, A3890401) and orange fluorescent beads (Fisher Scientific, F8800; Invitrogen, F8800) as previously described^19,20,56^. 10 µm thick sections of brain, duodenum, skin, and lung were cut from each tissue block on a cryostat (Leica, CM1860) prechilled to −20 °C using disposable blades (Leica, 14035838926), and the sections were collected onto the surface of coverslips also prechilled to −20 °C. The sections were gently melted onto the coverslips by briefly touching the opposite side of the coverslip with a gloved fingertip and were immediately refrozen. To enhance mechanical adhesion, the sections were allowed to dry inside the cryostat for 2 hours at −20 °C. Following drying, the sections were fixed with 4% (w/v) PFA in 1× PBS for either 20 minutes at 4 °C in a cold room or 1 hour at room temperature. The tissues were subsequently rinsed twice with 1× PBS for 2 minutes per wash at either 4 °C or room temperature. For permeabilization, tissues were washed with 70% ethanol and then transferred to a new 60 mm Petri dish with fresh 70% ethanol and stored at 4 °C overnight. Samples were stored for no longer than 7 days before proceeding to subsequent steps.

Tissues were prepared for MERFISH imaging following previously described protocols^20,25,26,56^. Briefly, an encoding probe hybridization solution consisting of 30% (v/v) formamide (Fisher Scientific, AM9342), 1 mg/mL yeast tRNA (Thermo, 15401029), and 10% (w/v) dextran sulfate (Sigma, S4030) in 2× saline-sodium citrate (SSC; Thermo, AM9765) was supplemented with 2 µM anchor probe (/5Acryd/TTG AGT GGA TGG AGT GTA ATT+ TT+ TT+ TT+ TT+ TT+ TT+ TT+ TT+ TT+ T, where /5Acryd/ denotes an acrydite modification and T+ indicates a locked nucleic acid; Integrated DNA Technologies) and the MERFISH probe library (35 µM for the 940-gene library, 25 µM for the 222-gene library, and 35 µM for the 449-gene library). Coverslips were removed from 70% ethanol and rinsed twice with 30% (v/v) formamide in 2× SSC for 5 minutes each at room temperature, then hybridized on a parafilm-lined 150 mm Petri dish with a 70 µL droplet of the encoding probe solution described above. To minimize evaporation of the hybridization solution, a moistened Kimwipe (Fisher, 06-666) was placed in the staining chamber, which was then incubated at 37 °C in a humidified incubator.

We leveraged multiple mouse libraries in this work: a 940-gene library introduced here, a 940-gene library published previously (940-gene [Cell 2024]), a 449-gene library, and a 222-gene library. We adjusted the incubation time to reflect differences in the per-probe concentration between these different libraries. We used 72 hours for the 940-gene, 940-gene (Cell 2024), and 449-gene libraries, and we used 48 hours for the 222-gene library.

Following hybridization, coverslips were carefully transferred to a 60 mm Petri dish and washed twice in 30% (v/v) formamide in 2× SSC for 30 minutes at 47 °C. Samples were then rinsed in a hydrogel solution composed of 4% (w/v) 19:1 acrylamide/bis-acrylamide (Bio-Rad, 1610144), 300 mM NaCl (Thermo, AM9759), 0.03% (w/v) ammonium persulfate (Sigma, 215589), and 0.15% (v/v) TEMED (Sigma, T7024) in 50 mM Tris-HCl pH 7 (Thermo, 15568-025). A glass plate (Gorilla Scientific, 6101) was cleaned and coated with Gel-Slick (Lonza, 50640), and the coverslip was inverted, sample-side down, onto a 60-100 µL droplet of gel solution on the glass plate. Excess gel solution was removed by gently pressing on the edges of the coverslip. The gel was allowed to polymerize for 2 hours at room temperature, after which the coverslip with the attached gel was gently separated from the glass plate using a razor blade and incubated in a digestion buffer consisting of 1:100 proteinase K (NEB, P8107S) in 0.25% (v/v) Triton X-100 (Sigma, T8787), 2% (v/v) sodium dodecyl sulfate (SDS; Thermo, AM9823) in 2× SSC. Samples were incubated in this digestion buffer at 37 °C for 2 days, with fresh buffer exchanged after the first day. The digestion buffer was subsequently removed by washing the samples in 2× SSC at room temperature, with six washes of 30 minutes each, to ensure thorough removal of residual SDS. After washing, samples were stored in 2× SSC at 4 °C for no more than one week before imaging.

### Fixed frozen sample preparation for MERFISH

Fixed frozen samples were sectioned, stained, and cleared using the same protocols as described for the fresh frozen MERFISH samples with the following notable differences. Because heavily fixed brain samples (48-hour fixation) showed poor adherence to coverslips, a few droplets of pre-chilled 70% ethanol were added to the coverslip and sections were transferred directly onto this droplet. As the droplet evaporated during the subsequent room temperature air-dry step, it would stretch and flatten the tissue, improving coverslip adherence. Aside from this modification, all samples were processed identically for subsequent steps. First, sections were briefly melted and refrozen on coverslips, then air-dried for 30-60 minutes at room temperature to further improve tissue adherence. Second, we stained samples with probes in a previously described two-step ‘post-gel’ protocol, designed to enhance the encoding probe penetration into samples that had been heavily fixed^56^. Briefly, samples were stained as above with encoding probe hybridization solution; however, this solution contained only 2 µM anchor probes. It did not contain encoding probes. Some samples were stained for 48 hours; however, as we found that 24 hours were sufficient for ample anchor probe staining, some samples were stained for this reduced duration.

The samples were then washed, gel-embedded, and cleared, as described above. To stain the samples with encoding probes, the gel was trimmed to just the region containing the sample of interest, and a hydrophobic barrier was drawn around the gel with a hydrophobic pen (Cosmo Bio, DAI-APAP-M-5) to limit the spread of encoding probe hybridization solution. 70 µL of encoding probe solution supplemented with encoding probes (35 µM for the 940-gene, 940-gene [Cell 2024], 737-gene, or 449-gene libraries and 25 µM for the 222-gene library) was then gently placed on the sample. Samples were then hybridized and washed as described for the fresh-frozen protocol using the same hybridization incubation durations. Samples were stored in 2× SSC at 4 °C until imaging.

### Fixed-frozen dual-gel sample preparation

We found that heavily fixed tissues would occasionally have degraded coverslip adherence and that during the sample staining and wash process small portions of the gel and sample would lift from the coverslip. This issue was more prominent with samples fixed in the cesium sulfate preservation buffer. To address this issue, we developed a dual-gel protocol to enhance the mechanical adherence of the sample. Immediately after the air-drying step post sectioning, slices were embedded in a thin polyacrylamide hydrogel using the same gel-embedding protocol described above. The samples were not cleared. After gelation, this first gel was trimmed to reduce excess gel, and the sample was processed using identical protocols for anchor-probe staining. The sample was then embedded in a second hydrogel, using the protocol described above, to covalently bond the anchor probes to this second gel. All subsequent steps were performed as described above for fixed frozen samples.

### Paraffin embedding and deparaffinization

To embed samples in paraffin, tissues were collected using one of the fixed protocols described above. To remove residual fixation buffer, samples were washed in water for 5 minutes at room temperature for a total of 5 washes. Samples were then dehydrated through a graded ethanol series consisting of 50 mL of 80%, 90%, and 95% ethanol for one hour each at 4 °C. The samples were then washed in 50 mL of 100% ethanol at 4 °C for a total of four times. Dehydrated tissue samples were then submitted to the Harvard Medical School Rodent Histopathology Core for standard paraffin processing and embedding. The core incubated samples in xylene (VWR, 89370-088) for 1.5 hours at room temperature for a total of two times, infiltrated tissues with paraffin (Leica, 3801340) for ∼4 hours at 60 °C, and embedded them in a final paraffin block (Leica, 3801320).

5 μm thick tissue sections were then mounted on coverslips prepared as described above. The paraffin was removed by placing coverslips in a glass Petri dish and incubating at 60 °C for 30 minutes, rapidly transferring the samples to a chemical fume hood, and then treating them with 3–5 mL of xylene. Samples were washed four times in xylene at room temperature, with 150 seconds of incubation for each wash. 3–5 mL of 100% ethanol was then added to the sample, and the sample was incubated for 3 minutes at room temperature. This ethanol wash was performed twice. The sample was then rehydrated by sequential incubation in 3-5 mL of 95% and 70% ethanol, with a 1-minute incubation in each. After rehydration, samples were immediately processed according to the fixed-frozen protocol as described above.

### Bulk RNA isolation for RNA integrity measures and bulk RNA sequencing

To extract RNA for either RNA integrity measurements or sequencing, tissues were dissected as described above and immediately submerged into either 1× PBS, commercial alcohol-based fixative PAXgene (Qiagen, 765312), cesium sulfate preservation buffer, homemade ammonium sulfate preservation buffer, RNAlater, other cesium-based solutions at varying temperatures, pHs, or sulfate concentrations as detailed for each experiment. Where necessary, the concentration of the preservation solutions was modulated by dilution of stock solutions with water. Samples were then either processed rapidly or allowed to incubate in the designated solution for the duration described. For PAXgene preservation, ileal samples were quickly dissected and stored in 2 mL of PAXgene Tissue FIX solution overnight. The PAXgene-fixed samples were then either directly processed for RNA isolation or washed in PBS for 1 hour at room temperature prior to RNA isolation.

To extract RNA, tissue samples were processed with the Direct-zol RNA Miniprep kit (Zymo, R2072). Briefly, harvested tissue was removed from the designated solution using forceps and excess solution was removed with a Kimwipe. The tissue was then placed into 700 µL of TRIzol (Invitrogen, 15596026) prechilled to 4 °C and immediately homogenized (Cole-Parmer, EW-04727-01) twice for 3 to 5 seconds each with samples held on an ice-water slurry. The lysate was vortexed and centrifuged at 14,000×g for 30 seconds, and 600 µL of the supernatant was transferred to a new 2 mL tube. An equal volume (600 µL) of ice-cold 100% ethanol was added and the mixture was vortexed. 750 µL of the mixture was transferred to a Spin IIICG column (Zymo, C1006) placed in a collection tube (Zymo, C1001) and centrifuged at 14,000×g for 30 seconds. The column was transferred to a new collection tube and the flow-through was discarded. For on-column DNase digestion, the column was first washed with 650 µL RNA Wash Buffer at 14,000×g for 30 seconds and then incubated for 15 minutes at room temperature with 80 µL of DNase I solution containing 5 µL DNase I (6 U/µL) and 75 µL DNA Digestion Buffer. The column was subsequently washed twice with 650 µL Direct-zol RNA PreWash Buffer at 14,000×g for 30 seconds each, followed by a final wash with 700 µL of RNA Wash Buffer at 14,000×g for 1 minute. The empty column was centrifuged for an additional 30 seconds at 14,000×g to remove residual buffer. RNA was eluted by transferring the column to a 1.5 mL DNA LoBind Tube (Fisher, 13-698-791) and adding 50 µL of nuclease-free water directly to the column matrix. After a 2-minute incubation at room temperature, the column was centrifuged at 14,000×g for 30 seconds to collect RNA. RNA concentration was measured using UV absorbance (Thermo, NanoDrop).

### RNA integrity measurements

As we would anticipate that MERFISH would be sensitive to the fragmentation of RNA, we used the RNA integrity number (RIN)—a measure of the degree to which RNA is intact—as a metric of RNA fragmentation. Briefly, bulk RNA extracted as described above was run on a microfluidic electrophoresis system (Agilent, TapeStation) using reagents and the instructions provided by the manufacturer (Agilent, RNA ScreenTape or High Sensitivity RNA ScreenTape). The equivalent RIN (eRIN) was provided by the software associated with this system.

To define how RNA integrity degrades with time differently between different tissues, we collected and sectioned fresh frozen blocks of mouse brain, lung, skin, and duodenum as described below. 5 to 10 slices of 20 µm-thick sections were collected and placed into 1.5 mL DNA LoBind tubes that were prechilled to −20 °C. To define the rapidity with which RNA can be degraded by the sample itself, we warmed these tubes through brief touch and then placed them at room temperature for various incubation times. Once the room temperature incubation was complete, we rapidly inhibited any degradation of the RNA by adding 800 µL of TRIzol. RNA was then extracted and RIN determined from these slices using the protocols described above.

### Bulk RNA-sequencing

Library preparation and sequencing of bulk RNA was performed by Novogene. Briefly, polyA mRNA was enriched via oligo(dT) beads (New England Biolabs, S1419), a sequencing library was created using a commercial library preparation kit (New England Biolabs, E7770), and then ∼20 million 150-bp paired-end sequences per sample were collected on an Illumina NovaSeq X Plus.

Raw sequencing reads were preprocessed using TrimGalore (https://www.bioinformatics.babraham.ac.uk/projects/trim_galore/) to remove adapter sequences and low-quality reads using default parameters. Paired-end sequencing reads were aligned to the mouse genome (GRCm39) using HISAT2^70^ and the standard analysis parameters.

Gene-level read counts were quantified using featureCounts^73^ Raw counts were normalized using the trimmed mean of M values (TMM) method implemented in the edgeR R package^74^. Log_2_-transformed TMM were then used to calculate Pearson correlation coefficients.

### RNase activity assay on tissue sections

To both support the notion that RNA is degraded by endogenous RNases within the tissue slice and provide a quantitative method to judge the degree to which RNase activity has been inhibited on individual slices collected from different tissues and preservation methods, we developed an on-slice RNase activity assay. Briefly, this assay is based on the release of a fluorescence quencher molecule linked to a fluorescent dye by a short RNA construct such that cleavage of the RNA releases the quencher, allowing fluorescence. Specifically, we used a commercially available construct for this purpose (RNaseAlert QC System V2; Thermo, 4479769).

We ran this assay in two formats: a per-slice format that allows for measurement on slices prepared using the same protocols for imaging and a per-well format that allows more rapid profiling of conditions and time points. In both formats, tissue blocks were prepared and 10 µm thick slices were collected as described above.

In the per-slice format, sections were air dried for 30 minutes at room temperature to increase tissue adherence to the coverslip, and a small region surrounding the tissue was created with a hydrophobic pen. The RNaseAlert substrate was diluted 10-fold into 1× RNaseAlert buffer in 1× PBS. Depending on section size, 40-200 µL of this solution was gently spread on each tissue section, and the sample was incubated at room temperature in the dark for various times. To image RNase activity, coverslips were placed in a gel imaging system (Axygen, Gel Documentation System-BL), illuminated with blue light, and images were acquired with a 250 ms exposure either immediately or after 5, 10, or 15 minutes of incubation (extended to 30 minutes when necessary). To quantify the increase in RNase activity, the average signal intensity within tissue regions, defined in ImageJ, was quantified at each time point. The reported values represent the difference between the 15-minute and immediate measurements, except for the mouse TMA, where values were calculated by subtracting the immediate measurement from the 30-minute incubation.

In the per-well format, sections were placed in individual microwells of a 96-well plate (Thermo Scientific, AB2396) and covered with 50 µL of the RNaseAlert solution described above. The plate was then placed on a qPCR machine (BioRad, CFX Opus 96 Real-Time PCR System), held at 37 °C, and imaged with the SYBR green channel every 1 minute for 2 hours.

### *In vitro* RNase A inactivation assay

To determine the efficacy of different approaches to RNase inactivation, we developed a simple *in vitro* RNase activity assay based on the RNaseAlert kit and RNase A. Briefly, 60 µL of 200 µg/mL RNase A, diluted from 20 mg/mL stock (Invitrogen, 12091021), was mixed with 940 µL of the solution to be tested. The sample was incubated at room temperature for 20 minutes, and then the solution was removed from the RNase A via size exclusion. Specifically, a size exclusion column (Amicon Ultra 3 kDa molecular weight cut off; Sigma, UFC5003) was prerinsed by spinning 500 µL of MilliQ H_2_O across the column at 14,000×g for 30 minutes. 475 µL of the sample was loaded into the column and spun at either 10,000×g for 20 minutes (cesium sulfate-based solution) or 14,000×g for 30 minutes (PBS-based, ammonium acetate-based, or RNAlater-based solution) at room temperature. 490 µL of 1×PBS was added to the column, and the same spin was performed again. This 1×PBS wash was performed for a total of two times, and then the remaining RNase A and solution was recovered by inverting the filter in a clean microcentrifuge tube and centrifuging at 1,000×g for 2 minutes. The recovered protein concentration was quantified via spectroscopy (NanoDrop).

To quantify the activity of the treated RNase A, the recovered RNase A was diluted to a standard concentration of 1 µg/mL in 1× PBS. 2.5 µL of this diluted RNase A was added to 47.5 µL of RNaseAlert reagent to bring the final working concentration to a 1:10 dilution of the RNaseAlert substrate and 1× RNaseAlert buffer. This mixture was placed in a micro-well of a 96-well plate, and the activity of the RNase A was measured as described above for the per-well format protocol.

### RNase activity assay on paraffin tissue microarray

Tissue microarray samples prepared as formalin-fixed paraffin-embedded tissues were purchased from Novus Biologicals (Fisher, NBP230225). The slides were deparaffinized as described above and then air dried at room temperature for 30 minutes. These samples were then processed using the per-slice format of the RNase activity assay as described above.

### MERFISH library design and construction

To characterize the samples described here, four mouse MERFISH libraries (940-gene, 940-gene [Cell 2024], 449-gene, or 222-gene) and one human MERFISH library (737-gene) were used. Encoding probes for each library have either been reported previously^75^ (940-gene [Cell 2024] library) or were designed here using previously described probe design methods^18,25^ (github.com/ZhuangLab/MERFISH_analysis).

The target regions of each library—the portion of the encoding probes that binds to RNAs—were designed using the same criteria. Briefly, 30-nt target regions for each gene were designed to have a GC fraction between 0.4 and 0.6, predicted melting temperature between 65 and 75 °C, and 20-nt overlap between target regions of the same gene. Target regions were selected that had a gene specificity and an isoform specificity greater than 0.7, where these specificity values were defined as described previously^18,25^. In some cases, the degree of homology between isoforms was too great to allow isoform distinction; thus, the isoform specificity requirement was removed. We aimed for 72 target regions for each targeted RNA but used all possible target regions for RNAs that were too short to support this number.

We generated constant Hamming Weight 4, minimum Hamming Distance 4, binary barcodes of sufficient length to cover the number of desired targets for each library while leaving some barcodes unassigned (so called ‘Blank’ barcodes) to serve as false positive controls. A previously published set of readout probes^18,20^ were associated with each bit in each of the barcodes. For some libraries these readout probes were first screened against the tissue to eliminate those with higher non-specific binding, which can decrease the rate of false positives^75^. Of the libraries used here, this pre-screening step was only performed for the 940-gene and the 737-gene libraries introduced here (as opposed to the previously published^25^ 940-gene library used only for the colon data presented here). Encoding probes were designed such that they contained 3 of the 4 associated readout probes for the barcode assigned to the targeted RNA and such that the full set of encoding probes targeting a given RNA had roughly equal representation of all readouts assigned to that RNA. Template libraries were designed by concatenating primers necessary for amplification. All template libraries were ordered from Twist Biosciences, and all primers were ordered from Integrated DNA Technologies.

Encoding probe libraries were amplified from these templates as previously described^17,18,56^. Briefly, the encoding probe template library was amplified with limited-cycle PCR and purified via spin column (Zymo, D4004 and C1012-50) to produce DNA templates for *in vitro* transcription. These templates were then *in vitro* transcribed to produce large quantities of single-stranded RNA (New England Biolabs, E2050S). The RNA was purified with home-made solid phase reversible immobilization (SPRI) magnetic beads^56^. The RNA was reverse transcribed and RNA template removed with alkaline hydrolysis. The final encoding probes were purified via SPRI beads and, where needed, concentrated with ethanol precipitation. The reverse transcription primer incorporated a 3’ RNA base, allowing the reverse transcription primer to be removed during alkaline hydrolysis.

### MERFISH imaging system

MERFISH data collection was performed on a home-built microscope and fluidics system as previously described^25,26^. Briefly, the imaging setup centered on a Nikon Ti2 microscope, fitted with either a 60× CFI PlanApo Lambda oil-immersion objective or a 10× CFI PlanApo Lambda air objective (both from Nikon). Sample positioning and focusing were managed using a motorized XY stage (Marzhauser, SCAN IM 130×85; Zaber, ASR100B120B-T3A-K0066) in combination with a piezoelectric objective positioner (Mad City Labs, Nano-F200). Excitation was provided in five channels from a laser light engine (Lumencor, Celesta) coupled into the microscope through a custom epi-illumination assembly optimized to uniformly illuminate an approximately a 230-μm square region of the specimen. The illumination path passed through a penta-band excitation filter (Semrock, FF01-391/477/549/639/741-25) and was directed onto the sample by a matching penta-band dichroic mirror (Semrock, FF421/491/567/659/776-Di01-25x36). Fluorescence from the sample was collected and separated into two detection channels via a TwinCam image splitter (Cairn Research), equipped with a long-pass dichroic element (Chroma, T750LPXRXT-UF2) that isolated emission beyond 750 nm from shorter wavelengths. Each channel was recorded by an independent CMOS camera (Hamamatsu, ORCA-Flash 4.0). The optical configuration provided an effective pixel size of roughly 105 nm at 60× magnification. Focus stability was maintained by a custom-built autofocus system, as described previously^25^. Samples were housed within a Bioptechs FCS2 flow chamber, where buffer exchange was performed using either a peristaltic (Gilson, MP1) or syringe (Hamilton, PSD4) pump. Fluid routing was governed by a series of computer-controlled valves (Hamilton; MVP4 with HVXM 8-5 valves), linked through a custom manifold and sipper assembly. All microscope operations, stage control, and fluid-handling routines were coordinated using custom software available at github.com/ZhuangLab/storm-control.

### MERFISH imaging

The sequential rounds of readout hybridization and imaging required for MERFISH were performed as described previously^17^. Repetitive staining of the sample leveraged four buffers. Readout hybridization buffer was comprised of 2× SSC buffer supplemented with 10% (v/v) ethylene carbonate (Alfa Aesar, A15735-36) and 0.25% (v/v) Triton X-100 (Sigma, T8787) and 15 nM each of the appropriate pair of readout probes. Readout wash buffer was identical to the readout hybridization buffer but did not contain readout probes. Cleavage buffer was comprised of 50 mM tris(2-carboxyethyl)phosphine (TCEP; GoldBio, TCEP25) in 2× SSC. Imaging buffer was composed of 300 mM NaCl, 50 mM Tris-HCl, 5 mM EDTA, 50 µM Trolox-quinone (prepared as described previously^56,76^), 0.5 mg/mL Trolox (Abcam, ab120747), 0.2% (v/v) rPCO (OYCAmericas, 46852004), and 5 mM protocatechuic acid (Sigma, 37580).

Before loading the sample on the microscope, the first two readout probes were stained by hybridizing the sample in 3 mL of the readout hybridization buffer for 15 minutes at room temperature, and the sample was then washed in 3 mL of readout wash buffer, supplemented with 1 mg/mL DAPI (Fisher Scientific, D1306), for 10 minutes at room temperature. The sample was then washed in 2× SSC. After washing, the sample was immediately loaded on the microscope.

Each repetitive staining and imaging round comprised the following steps. The sample was washed with ∼1-2 mL of imaging buffer, then the desired fields-of-view (FOV) were imaged in all appropriate color channels. For all samples, the full volume of the slice was imaged by collecting seven z-planes, spaced by 1.5 µm, and a single z-plane was captured using the 535-nm channel to locate fiducial beads on the coverslip surface. In the first imaging round, the 405-nm channel was used to image nuclei via the DAPI stain. After imaging, the fluorophores were cleaved from the readout probe by flowing ∼3 mL of cleavage buffer across the sample and incubating for 15 minutes at room temperature. Excess TCEP was then removed by flowing ∼3 mL of 2× SSC. Then ∼3 mL of readout hybridization buffer was flown across the sample, and it was incubated in this solution for 15 minutes at room temperature to stain the readout probes for the next round. Excess readout probes were then washed from the sample by flowing ∼3 mL of readout wash buffer across the sample. This process was then repeated as needed until the entire barcodes were measured.

### Hematoxylin and eosin (H&E) imaging

5 µm thick tissue sections prepared from FFPE blocks, as described above, were mounted on coverslips and submitted to the Harvard Medical School Rodent Histopathology Core for standard H&E staining. The stained sections were then imaged using a Nikon Ti-2 microscope under brightfield illumination equipped with a FLIR Blackfly S BFS-U3-122S6C camera.

## QUANTIFICATION AND STATISTICAL ANALYSIS

### MERFISH image decoding

MERFISH images were processed with a previously established pipeline^19,20^, available at (github.com/ZhuangLab/MERFISH_analysis). In short, images from separate rounds were spatially warped based on a set of orange fiducial beads imaged in each round, optimized to normalize fluorescent intensity between imaging rounds, and individual RNAs identified by a pixel-based soft decoding approach that matches pixel traces between imaging rounds to the barcoding scheme and aggregates adjacent pixels assigned to the same barcode to a single RNA molecule. Detected RNA were first filtered based on the number of adjacent pixels aggregated (the area) and the L2-normalized intensity across all rounds (the brightness).

### Cell segmentation

RNA were segmented into cells with Baysor^77^ (version 0.7.0 for all samples, except for the colonic Swiss roll, which was processed with version 0.4.3). The Baysor parameters were tuned for every tissue type to reflect differences in cell size and RNA density between tissues and libraries. However, within a tissue-library combination, the Baysor parameters were constant between different preparation methods. In all cases, 3D segmentation was performed except for the colonic Swiss roll which was 2D. For lung, duodenum, skin and intestinal biopsies, segmentation was performed without blank barcodes. For these datasets, after Baysor segmentation, blank barcodes were assigned to cells if they fell within 2 µm of a non-blank RNA molecule assigned to that cell.

### Cell annotation and single-cell analysis

To provide a joint labeling of cell populations collected within each of the mouse tissues used for our preparation method screening—lung, duodenum, skin, and brain—we leveraged a unified label transfer approach. By leveraging a unified reference per tissue, we were also able to define a quantitative measure of the relative degree to which different data quality can be used to resolve different cell populations. To build the needed single-cell references, we turned to published single-cell RNA sequencing (scRNA-seq) data where available^49–54^.

For the lung, we downloaded published data^51^ from the lungmap portal (https://www.lungmap.net/dataset/?dataset_id=LMEX0000004397). All cells within the file (LungMAP_MouseLung_CellRef.v1.1.h5ad) were used here. These data provided multiple tiers of cell annotation, of which we transferred the 40 labels in tier 3 annotation (labelled celltype_level3) onto our data. For ease of visualization, we consolidated these into 28 annotations.

For the duodenum, no single reference dataset encompassed all major cellular compartments; thus, we combined three complementary public datasets representing the epithelial^52^, stromal^53^, and immune^54^ compartments of the mouse small intestine. Epithelial data was downloaded from https://cellxgene.cziscience.com/collections/3db5617e-9f12-4eb4-8416-94893a0d7c46, stromal data from GEO (GSM5469262), and immune data from GEO (GSE221006). The expression matrices associated with these cells were concatenated, and the published cell annotations were used.

For the brain, we turned to a recent whole-brain scRNA-seq^50^ and spatial atlas^49^, and we cut these data to represent cells collected from the region of the brain we characterized (roughly Bregma -1.07mm to 1.60 mm). Specifically, we downloaded data from https://allen-brain-cell-atlas.s3.us-west-2.amazonaws.com/index.html#expression_matrices/. We then restricted the datasets collected with either the v2 or v3 10X chemistries (labeled as WMB-10Xv2 or WMB-10Xv3) to brain regions extending into our slices, which were contained within one of the following region labels: CTXsp, HPF, HY, Isocortex-1, Isocortex-2, Isocortex-3, Isocortex-4, TH, PAL, and STR. As these regions extend well beyond the region we characterized in our slices and, thus, contain many cell populations not expected in our slices, we leveraged a recent MERFISH atlas^49^ to identify the cell type labels associated with the scRNA-seq reference found within our slices. We examined slices Zhuang-ABCA-1.072 to Zhuang-ABCA-1.083 and Zhuang-ABCA-2.034 to Zhuang-ABCA-2.038, which flank ∼50 µm around the slice we identified and kept all cell type labels that contained at least 30 cells within these slices. We further excluded two labels that were associated with cells in the olfactory bulb, leaving 128 cell population labels in total. Finally, 500,000 cells were randomly down-sampled for label transfer. In the course of this analysis, we identified one gene measured with MERFISH, *Ptgdr*, that was an outlier as compared to bulk RNA sequencing data and, thus, was likely influenced by false positives. We set this gene aside for all analysis. We also found that this gene enriched within populations of cells identified in the fresh frozen preparation, and we excluded all MERFISH cells associated with this population from further analysis.

All replicate MERFISH measurements for a given tissue and preparation condition were then jointly integrated with the appropriate reference using the default parameters associated with the Harmony^71^ algorithm as implemented in Scanpy^72^. To integrate the datasets, only genes present in both MERFISH and scRNA-seq were considered, count matrices were normalized to a fixed count per cell, log_10_-transformed, and z-scored. The number of principal components (PC) used for the integration was determined by a pseudo-JackStraw procedure as described previously^26^. Once integrated, scRNA-seq labels were transferred to each MERFISH cell by identifying the ten nearest scRNA-seq cells for each MERFISH cell in this PC space. The most frequent label was transferred.

Owing to differences in developmental stage, anatomical location, and the relatively limited number of cells, we decided not to utilize the scRNA-seq atlas from Joost et al.^55^ for cell-label transfer in the skin. Instead, we annotated the MERFISH data produced by the highest quality preparation (fixation at room temperature in the cesium sulfate preservation buffer) and then used the above protocol to transfer these annotations to all other skin MERFISH data collected with different preparation methods. MERFISH data were normalized, log_10_-transformed, and z-scored as above. PCs were calculated and a reduced set selected, as above, and Leiden clustering was performed on a nearest neighbor graph using the default settings in Scanpy. Cell type labels were assigned using established marker gene expression.

To assess the biological resolution of a given MERFISH data set, we defined a confidence score for the transfer of labels. Specifically, as we sought to define the inherent resolution of the MERFISH data itself, independent of the biological resolution provided by the reference scRNA-seq data, we reanalyzed the replicates for each MERFISH preparation method on each tissue using the protocols as described above, explicitly excluding the reference data. In the reduced PC space defined by this analysis, we identified the 10 nearest MERFISH cells to each MERFISH cell and computed the confidence of the transferred label from the number of these neighbors that shared the same label. In this sense, this confidence score can be thought of as a measure of the coherence of the biological labels applied. Cells which are not inherently well defined by the MERFISH data will tend to be intermixed with other, often similar cell types, and, thus, will have a lower confidence score. In contrast, cell populations that are well defined would tend to co-occur with cells assigned the same label and, thus, have a higher confidence score. It is important to note that the quality of the MERFISH data alone is not the only factor that shapes this confidence score, as cell type abundance is also an important feature that sets the fraction of cells within a given neighborhood assigned the same label. Thus, increasing confidence score within a given cell type as the preparation is varied is a good indication that the resolving power of MERFISH has been improved; however, caution should be applied in interpreting relative confidence scores between cell types.

For display purposes, we occasionally reduced the number of listed labels by combining biologically similar cell population labels.

