## Supplemental Information for "RNA-aware tissue preservation workflows for high-quality spatial transcriptomics"

### SUPPLEMENTAL FIGURES

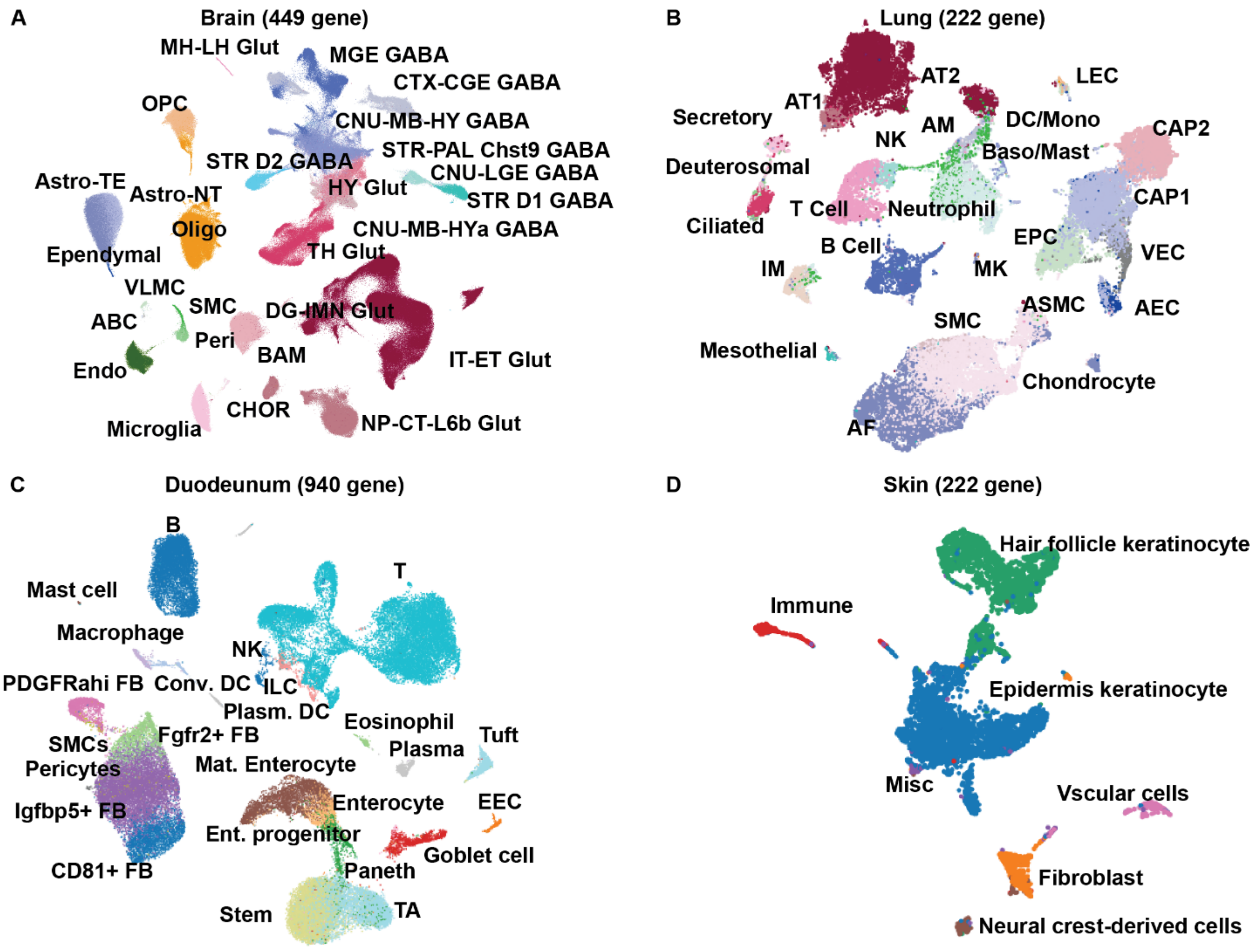


**Figure S1. MERFISH panels recapitulate the expected cell type diversity of the four profiled mouse tissues.**

(A-D) UMAP representation of the diversity of published brain (A), lung (B), duodenum (C), and skin (D) single-cell RNA sequencing data. Cells are colored by the published cell-type annotations with some cell types consolidated to increase visibility. The brain data were derived from Yao et al., including only 500,000 randomly subsampled cells relevant to the brain region profiled here, and were analyzed with only the genes in the 449-gene library. The lung data were derived from Guo et al. and were analyzed with only the genes in the 222-gene library. The duodenum data were derived from Zwick et al. (epithelial), Pærregaard et al. (stromal), and Wang et al. (immune) and were analyzed with only the genes in the 940-gene library introduced here. The skin data were taken from Joost et al. and were analyzed using only the genes in the 222-gene library.

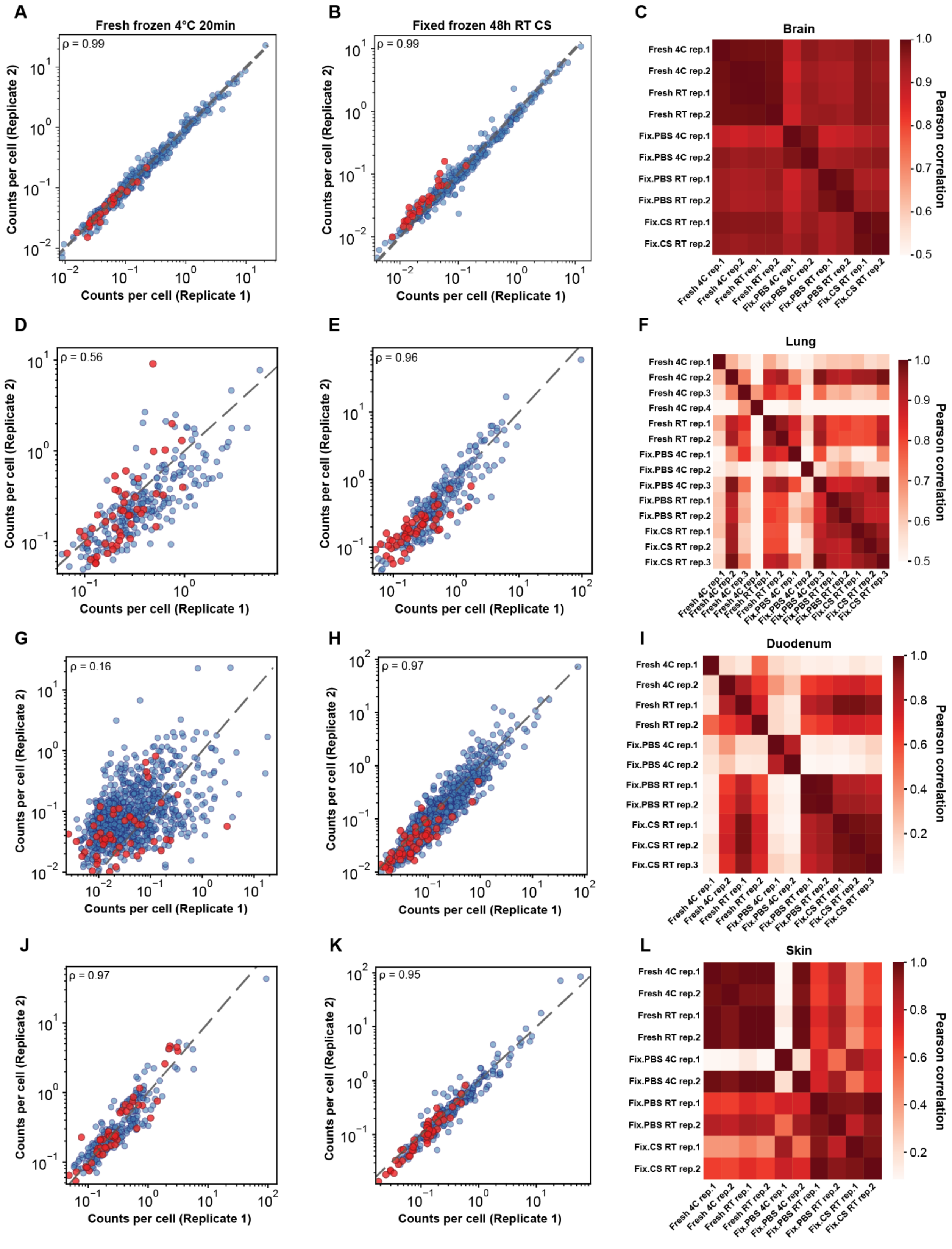


**Figure S2. Reproducibility of MERFISH measurements across different tissue types and sample preparation methods.**

(A) The average RNA abundance per cell for each gene determined via MERFISH for one biological replicate of the mouse brain prepared with a fresh frozen protocol that uses a 20-minute 4 °C post-fix versus that derived from a second biological replicate prepared with the same protocol. ρ represents the Pearson correlation coefficient of the $\log_{10}$ expression values. Blue markers represent RNAs while red markers represent barcodes not assigned to an RNA, i.e., ‘Blank’ barcodes.

(B) As in (A) but for two replicates of the mouse brain prepared with a fixed frozen protocol leveraging the cesium sulfate preservation buffer.

(C) The pairwise Pearson correlation coefficients for the $\log_{10}$ expression values for all biological replicates prepared with each of the five different preparation methods profiled for the mouse brain.

(D-F) As in (A)-(C) but for the mouse lung.

(G-I) As in (A)-(C) but for the mouse duodenum.

(J-L) As in (A)-(C) but for the mouse skin.

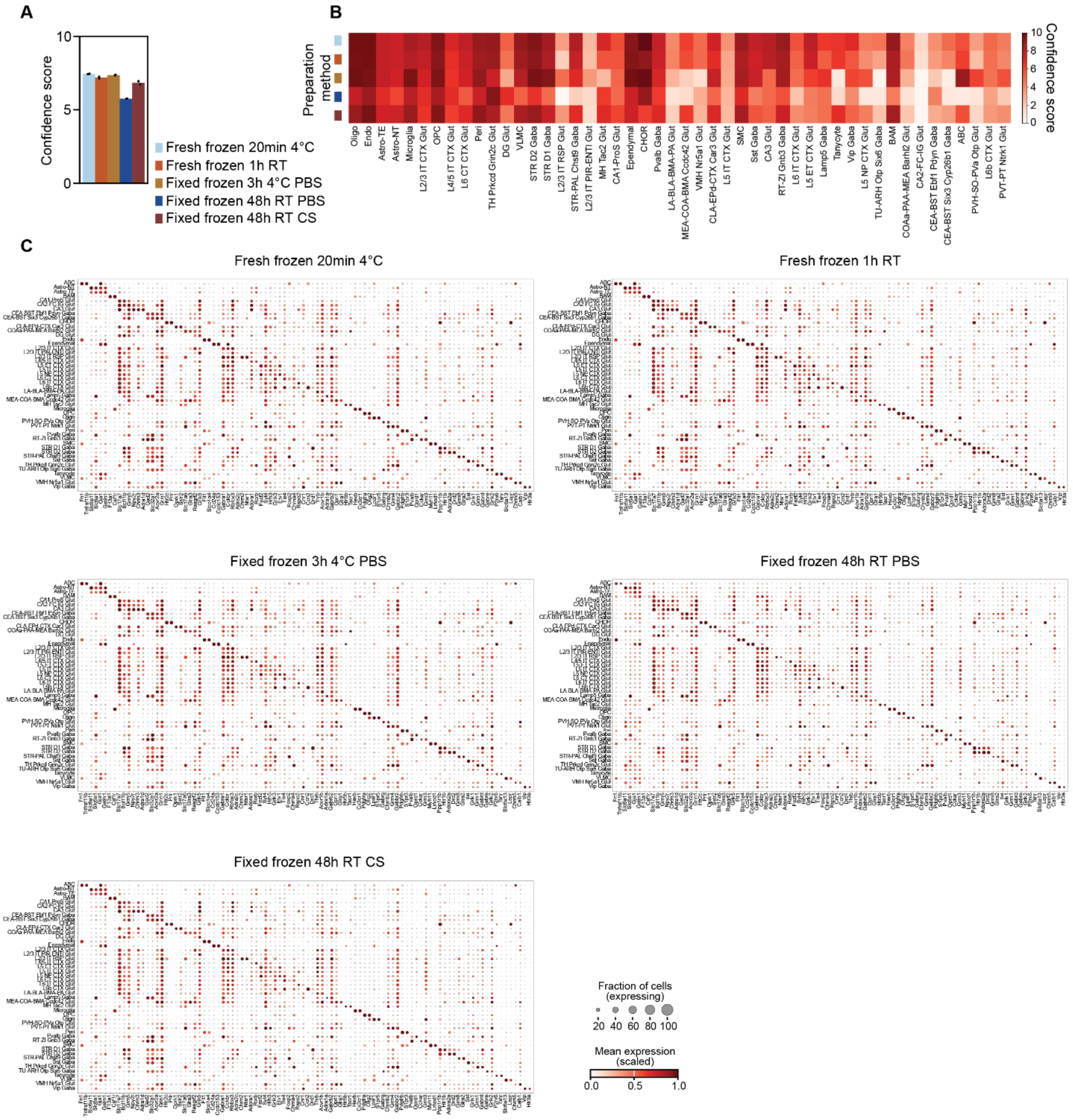


**Figure S3. The quality of cell type assignment for MERFISH data collected for the mouse brain under different preparation methods.**

(A) The average confidence score with which cell type labels can be assigned with MERFISH data alone. This score goes from 0 to 10 and is calculated by averaging the number of cells in a neighborhood of 10 cells that share the same cell type label as a central cell. The neighboring cells are determined by distance in a reduced gene expression space (Methods). A greater coherence in the cell type labels within the neighborhood of a given cell is indicative of a greater degree of confidence in the cell type label assigned to that cell. Bars represent the average confidence scores across two replicate MERFISH measurements indicated by individual black dots for the listed protocols.

(B) The confidence score, as defined in (A), but averaged across all cells assigned to each listed cell type label. Colored labels on rows represent the preparation method as indicated in (A).

(C) Expression of the top two differentially expressed genes for each listed cell type, with each gene shown only once across cell types (if a gene is identified in multiple cell types, it is retained for one and removed from the others). Marker size represents the fraction of cells expressing at least one copy of that gene, while marker color represents the average expression level normalized to the cell type with the highest expression of that gene. Individual panels represent the aggregate data from two biological replicates for each preparation method listed in the title.

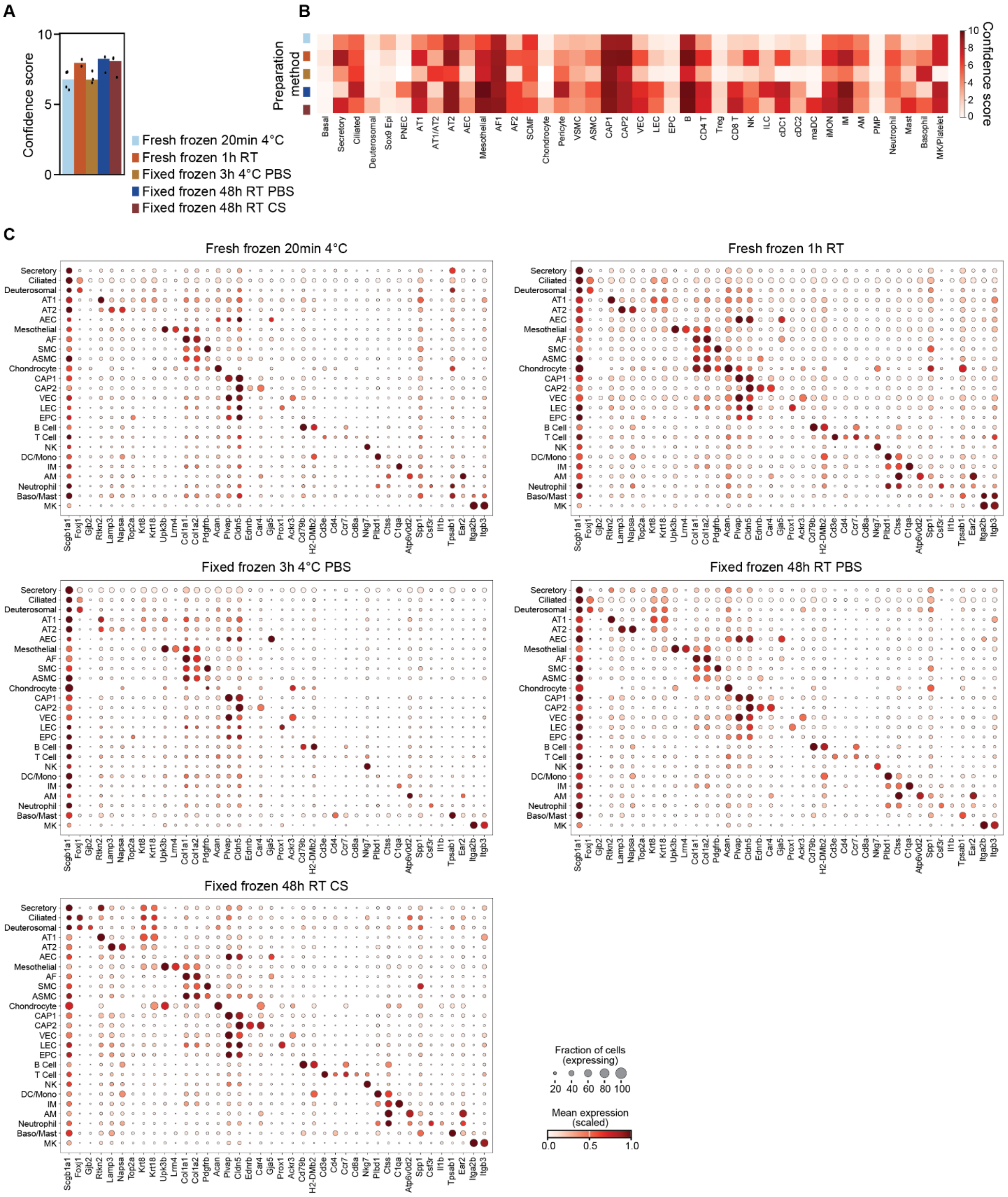


**Figure S4. The quality of cell type assignment for MERFISH data collected for the mouse lung under different preparation methods.**

(A) The average confidence score with which cell type labels can be assigned with MERFISH data alone. The confidence score is defined as in Figure S3 (Methods). Bars represent the average confidence scores across two replicate MERFISH measurements indicated by individual black dots for the listed protocols.

(B) The confidence score, as defined in (A), but averaged over all cells assigned to each listed cell type label. Colored labels on rows represent the preparation method as indicated in (A).

(C) The average marker gene expression within cells assigned the listed cell type labels. Marker size represents the fraction of cells expressing at least one copy of that gene, while marker color represents the average expression level normalized to the cell type with the highest expression of that gene. Individual panels represent the aggregate data from at least two biological replicates for the preparation method listed in the title.

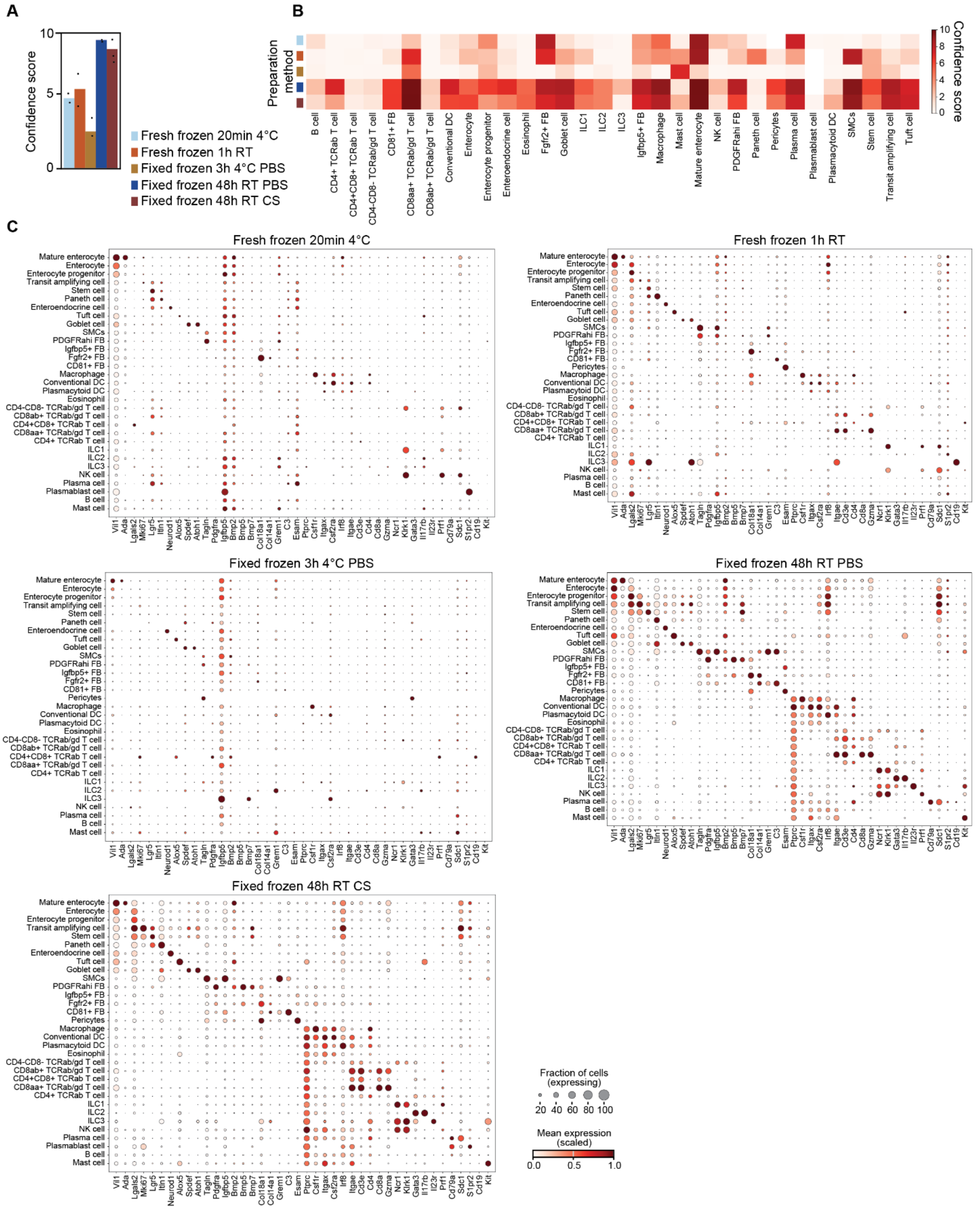


**Figure S5. The quality of cell type assignment for MERFISH data collected for the mouse duodenum under different preparation methods.**

(A) The average confidence score with which cell type labels can be assigned with MERFISH data alone. The confidence score is defined as in Figure S3 (Methods). Bars represent the average confidence scores across two replicate MERFISH measurements indicated by individual black dots for the listed protocols.

(B) The confidence score, as defined in (A), but averaged over all cells assigned to each listed cell type label. Colored labels on rows represent the preparation method as indicated in (A).

(C) The average marker gene expression within cells assigned the listed cell type labels. Marker size represents the fraction of cells expressing at least one copy of that gene, while marker color represents the average expression level normalized to the cell type with the highest expression of that gene. Individual panels represent the aggregate data from at least two biological replicates for the preparation method listed in the title.

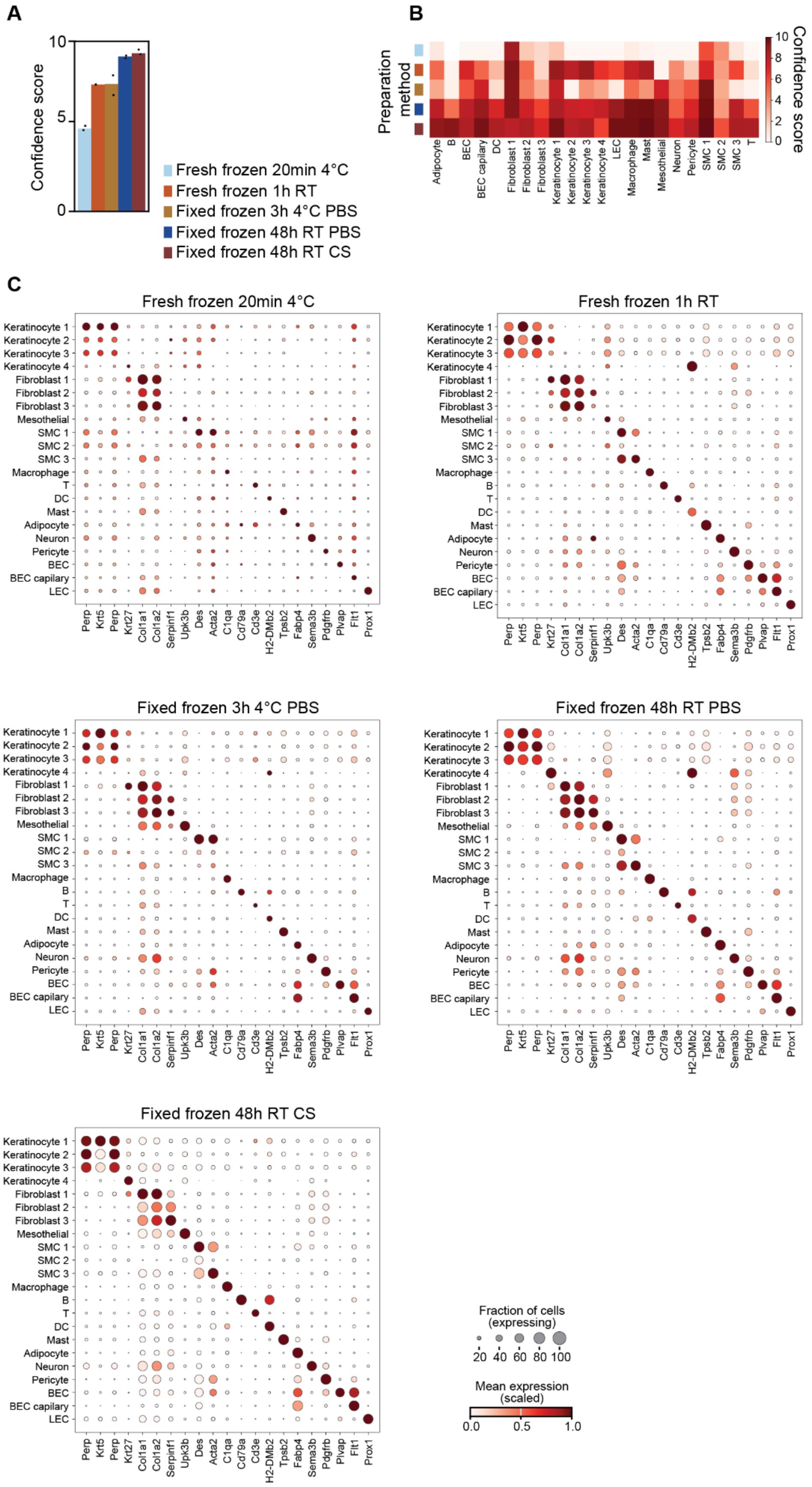
A
Confidence score
10
5
0
Fresh frozen 20min 4°C
Fresh frozen 1h RT
Fixed frozen 3h 4°C PBS
Fixed frozen 48h RT PBS
Fixed frozen 48h RT CS
B
Preparation method
Confidence score
Adipocyte
B
BEC
BEC capilary
DC
Fibroblast 1
Fibroblast 2
Fibroblast 3
Keratinocyte 1
Keratinocyte 2
Keratinocyte 3
Keratinocyte 4
LEC
Macrophage
Mast
Mesothelial
Neuron
Pericyte
SMC 1
SMC 2
SMC 3
T
C
Fresh frozen 20min 4°C
Fresh frozen 1h RT
Fixed frozen 3h 4°C PBS
Fixed frozen 48h RT PBS
Fixed frozen 48h RT CS
Perp
Krt5
Perp
Krt27
Col1a1
Col1a2
Serpinf1
Upk3b
Des
Acta2
C1qa
Cd79a
Cd3e
H2-DMb2
Tpsb2
Fabp4
Sema3b
Pdgfrb
Plvap
Flt1
Prox1
Fraction of cells (expressing)
20 40 60 80 100
Mean expression (scaled)
0.0 0.5 1.0

**Figure S6. The quality of cell type assignment for MERFISH data collected for the mouse skin under different preparation methods.**

(A) The average confidence score with which cell type labels can be assigned with MERFISH data alone. The confidence score is defined as in Figure S3 (Methods). Bars represent the average confidence scores across two replicate MERFISH measurements indicated by individual black dots for the listed protocols.

(B) The confidence score, as defined in (A), but averaged over all cells assigned to each listed cell type label. Colored labels on rows represent the preparation method as indicated in (A).

(C) The average marker gene expression within cells assigned the listed cell type labels. Marker size represents the fraction of cells expressing at least one copy of that gene, while marker color represents the average expression level normalized to the cell type with the highest expression of that gene. Individual panels represent the aggregate data from at least two biological replicates for the preparation method listed in the title.

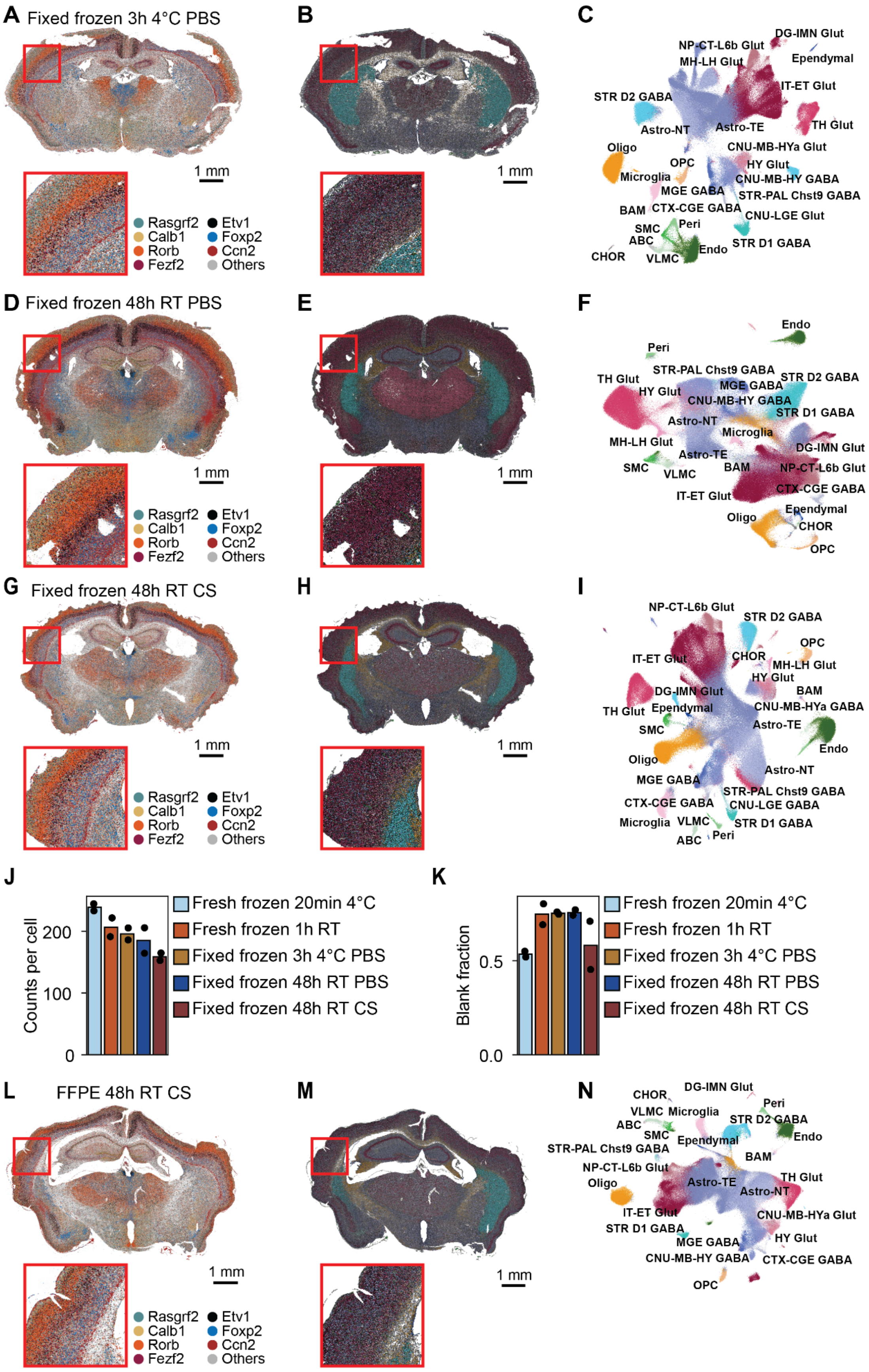

A Fixed frozen 3h 4°C PBS
B
C
1 mm
Rasgrf2
Calb1
Rorb
Fezf2
Etv1
Foxp2
Ccn2
Others
DG-IMN Glut
NP-CT-L6b Glut
MH-LH Glut
Ependymal
IT-ET Glut
STR D2 GABA
Astro-NT
Astro-TE
TH Glut
Oligo
CNU-MB-HYa Glut
OPC
HY Glut
Microglia
CNU-MB-HY GABA
MGE GABA
STR-PAL Chst9 GABA
BAM
CTX-CGE GABA
CNU-LGE Glut
SMC
Peri
ABC
Endo
STR D1 GABA
CHOR
VLMC
D Fixed frozen 48h RT PBS
E
F
G Fixed frozen 48h RT CS
H
I
J
Counts per cell
0
100
200
Fresh frozen 20min 4°C
Fresh frozen 1h RT
Fixed frozen 3h 4°C PBS
Fixed frozen 48h RT PBS
Fixed frozen 48h RT CS
K
Blank fraction
0.0
0.5
L FFPE 48h RT CS
M
N

**Figure S7. Fixed frozen protocols produce high-quality MERFISH in the mouse brain.**

(A and B) Spatial distribution of 7 of 449 mRNAs (A) or of the identified cells (B) measured with MERFISH in a coronal section of the mouse brain prepared with a fixed frozen protocol that leverages a 3-hour 4 °C fixation.

(C) UMAP representation of the cellular diversity seen in two biological replicates of the mouse brain with the protocol in (A), colored by major cell type labels transferred to the data from a reference dataset (Methods). The colors are the same as in (B).

(D-F) As in (A-C) but for MERFISH measurements of the mouse brain prepared with a fixed frozen protocol that leverages a 48-hour, room-temperature fixation.

(G-I) As in (A-C) but for a mouse brain prepared with a fixed frozen protocol that leverages a 48-hour, room-temperature fixation in a cesium sulfate preservation buffer.

(J and K) The average RNA counts per cell (J) or the average false positive counts per cell (K) observed for the mouse brain prepared with different preservation methods. Bars represent average over individual biological replicates represented as markers.

(L-N) As in (A-C) but for a mouse brain prepared with a paraffin embedded fixed protocol in which tissues were fixed for 48 hours at room temperature in the cesium sulfate preservation buffer.

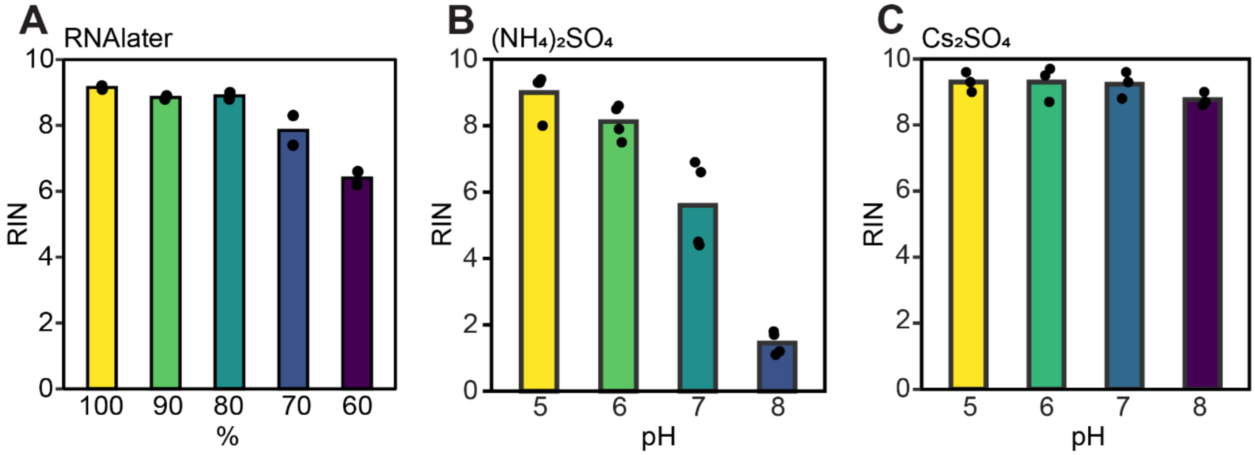


**Figure S8. Properties of RNA preservation with various sulfate solutions.**

(A) The RIN of RNA extracted from mouse duodenal samples stored in different dilutions of RNAlater into water for 48 hours at room temperature.

(B) The RIN of RNA extracted from mouse duodenal samples stored for 48 hours at room temperature in an ammonium sulfate preservation solution adjusted to have the listed pH.

(C) As in (B) but for samples stored in a cesium sulfate preservation solution adjusted to have the listed pH. Bars represent averages over individual biological replicates, which are denoted as markers.

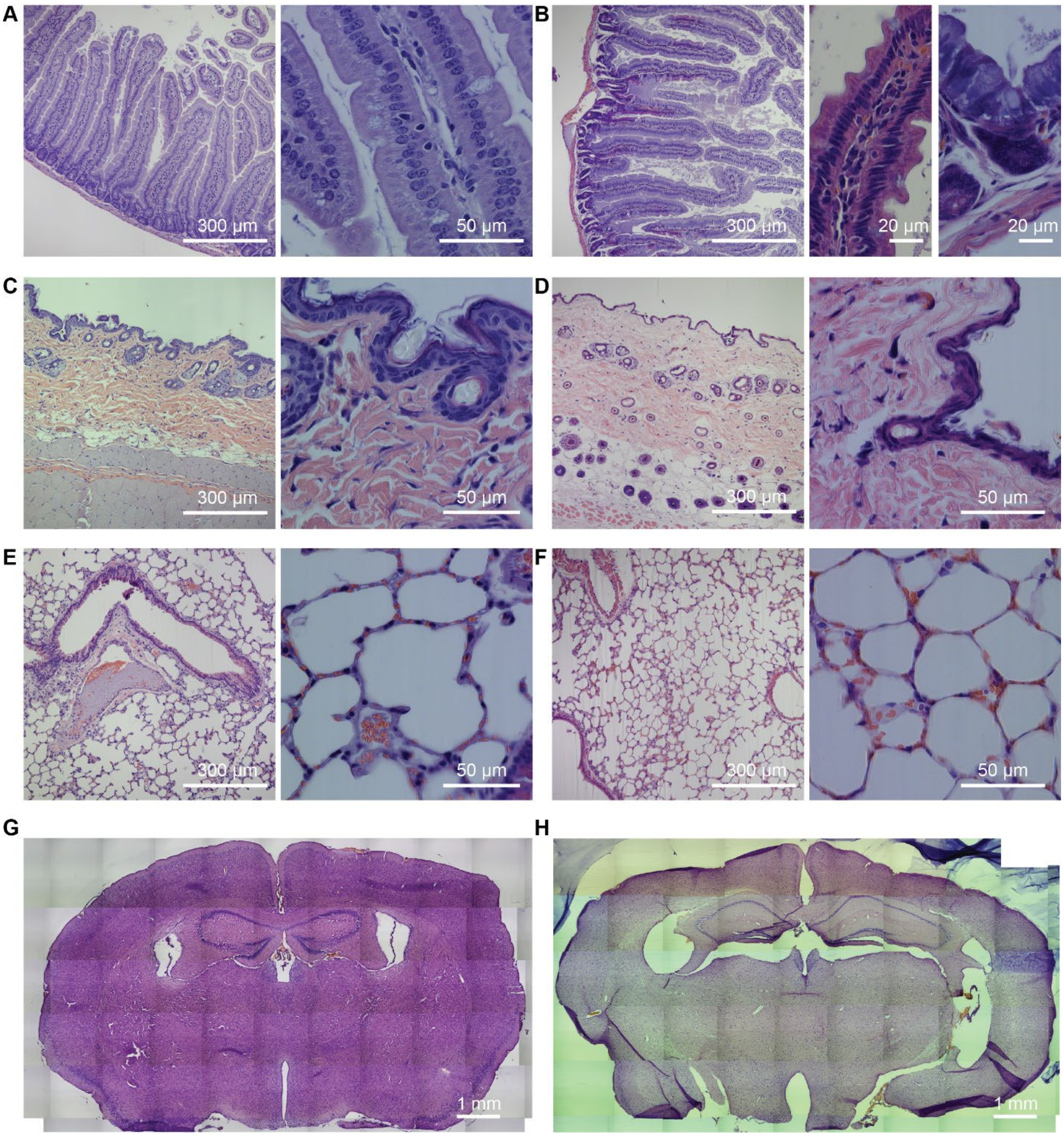


**Figure S9. Cesium-sulfate-based fixation produces some fixation artifacts while largely preserving the morphological structure of tissues.**

(A-H) Hematoxylin and eosin (H&E)–stained sections of duodenum (A and B), skin (C and D), lung (E and F), and brain (G and H) prepared with a paraffin embedded fixed protocol in which tissues were fixed for 48 hours at room temperature in PBS (A,C,E,G) or in the cesium sulfate preservation buffer (B,D,F,H).

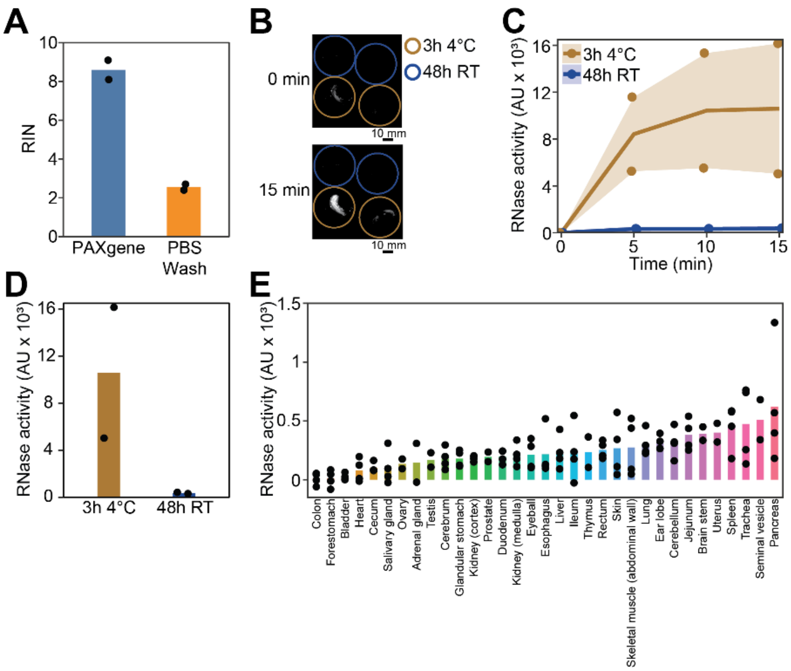


**Figure S10. Reversible alcohol fixation and residual RNase activity in paraffin-embedded fixed samples.**

(A) RIN measured for bulk RNA extracted from mouse ileum after storage in PAXgene overnight at room temperature or after a 60-min wash in PBS following this treatment (Methods). Each treatment has two biological replicates denoted by black dots.

(B) Images of a per-slice RNase activity measured by the generation of fluorescence from a test RNA in arbitrary units (Methods) for two replicate slices of paraffin-embedded mouse duodenal samples fixed either at 4 °C for 3 hours (brown circles) or at room temperature for 48 hours (blue circles).

(C) Integrated fluorescence over the tissue slices shown in (B) versus time at room temperature with the initial fluorescence subtracted from the initial time. Markers represent replicates and the line represents the average.

(D) Average RNase activity for the duodenal samples in (B) quantified as the measured fluorescence at 15 minutes minus that measured at 0 minutes. Markers represent replicates and bars represent the average.

(E) RNase activity measured as in (B) but for two replicates of a tissue microarray (TMA) that contained multiple biological replicates of formalin-fixed paraffin embedded cores of the listed mouse tissues. Here the reported activity represents the average fluorescence measured at 30 minutes minus the autofluorescence measured at 0 minutes. Bars represent averages and markers represent different cores from different mice. While detectable RNase activity was observed in many tissue cores, these results should not be used to compare relative RNase activity across tissues, as variable degrees of fixation were applied across tissue cores. Instead, these data are intended only to demonstrate that residual RNase activity is present in commercial TMA samples.
